# Predicting the Sequence-Dependent Backbone Dynamics of Intrinsically Disordered Proteins

**DOI:** 10.1101/2023.02.02.526886

**Authors:** Sanbo Qin, Huan-Xiang Zhou

**Affiliations:** Department of Chemistry, University of Illinois Chicago, Chicago, IL 60607, USA; Department of Physics, University of Illinois Chicago, Chicago, IL 60607, USA

**Keywords:** NMR spectroscopy, phase separation, intrinsically disordered proteins, backbone dynamics

## Abstract

How the sequences of intrinsically disordered proteins (IDPs) code for functions is still an enigma. Dynamics, in particular residue-specific dynamics, holds crucial clues. Enormous efforts have been spent to characterize residue-specific dynamics of IDPs, mainly through NMR spin relaxation experiments. Here we present a sequence-based method, SeqDYN, for predicting residue-specific backbone dynamics of IDPs. SeqDYN employs a mathematical model with 21 parameters: one is a correlation length and 20 are the contributions of the amino acids to slow dynamics. Training on a set of 45 IDPs reveals aromatic, Arg, and long-branched aliphatic amino acids as the most active in slow dynamics whereas Gly and short polar amino acids as the least active. SeqDYN predictions not only provide an accurate and insightful characterization of sequence-dependent IDP dynamics but may also serve as indicators in a host of biophysical processes, including the propensities of IDP sequences to undergo phase separation.

## Introduction

Intrinsically disordered proteins (IDPs) or regions (IDRs) do not have the luxury of a three-dimensional structure to help decipher the relationship between sequence and function. Instead, dynamics has emerged as a crucial link between sequence and function for IDPs (1). Nuclear magnetic resonance (NMR) spin relaxation is a uniquely powerful technique for characterizing IDP dynamics, capable of yielding residue-specific information (2). Backbone ^15^N relaxation experiments typically yield three parameters per residue: transverse relaxation rate (*R*_2_), longitudinal relaxation rate (*R*_1_), and steady-state heteronuclear Overhauser enhancement (NOE). While all three parameters depend on ps-ns dynamics, *R*_2_ is the one most affected by slower dynamics (10s of ns to 1 μs). An increase in either the timescale or the amplitude of slower dynamics results in higher *R*_2_. For IDPs, *R*_2_ is also the parameter that exhibits the strongest dependence on sequence (1, 2).

*R*_2_ was noted early on as an important indicator of residual structure in the unfolded state of the structured protein lysozyme (3). This property has since been measured for many IDPs to provide insight into various biophysical processes. Just as the residual structure in the unfolded state biases the folding pathway of lysozyme (3), a nascent α-helix in the free state of Sendai virus nucleoprotein C-terminal domain (Sev-NT), as indicated by highly elevated *R*_2_ (4), biases the coupled binding and folding pathway in the presence of its target phosphoprotein (5). Local secondary structure preformation also facilitates the binding of yes-associated protein (YAP) with its target transcription factor (6). Likewise a correlation has been found between *R*_2_ in the free state and the membrane binding propensity of synaptobrevin-2: residues with elevated *R*_2_ have increased propensity for membrane binding (7). *R*_2_ in the free state has also been used to uncover factors that promote liquid-liquid phase separation of IDPs. For example, a nascent α-helix (shown by elevated *R*_2_) is important for the phase separation of the TDP-43 low-complexity domain, as both the deletion of the helical region and a helix-breaking mutation (Ala to Pro) abrogates phase separation (8). Similarly, nascent α-helices in the free state of cytosolic abundant heat-soluble 8 (CAHS-8), upon raising concentration and lowering temperature stabilize to form the core of fibrous gels (9). For the hnRNPA1 low-complexity domain (A1-LCD), aromatic residues giving rise to local peaks in *R*_2_ also mediate phase separation (10).

Both NMR relaxation data and molecular dynamics (MD) simulations have revealed determinants of *R*_2_ for IDPs. It has been noted that the flexible Gly tends to lower *R*_2_ whereas secondary structure and contact formation tend to raise *R*_2_ (11). This conclusion agrees well with recent MD simulations (1, 12-14). These MD studies, using IDP-specific force fields, are able to predict *R*_2_ in quantitative agreement with NMR measurements, without ad hoc reweighting as done in earlier studies. According to MD, most contact clusters are formed by local sequences, within blocks of up to a dozen or so residues (1, 12, 14). Tertiary contacts can also form but are relatively rare; as such their accurate capture requires extremely extensive sampling and still poses a challenge for MD simulations. Contrary to Gly, aromatic residues have been noted as mediators of contact clusters (3, 10).

Schwalbe et al. (15) introduced a mathematical model to describe the *R*_2_ profile along the sequence for lysozyme in the unfolded state. The *R*_2_ value of a given residue was expressed as the sum of contributions from this residue and its neighbors. This model yields a mostly flat profile across the sequence, except for falloff at the termini, resulting in an overall bell shape. Klein-Seetharaman et al. (3) then fit peaks above this flat profile as a sum of Gaussians. Cho et al. (16) proposed bulkiness as a qualitative indicator of backbone dynamics. Recently Sekiyama et al. (17) calculated *R*_2_ as the geometric mean of “indices of local dynamics”; the latter were parameterized by fitting to the measured *R*_2_ for a single IDP. All these models merely describe the *R*_2_ profile of a given IDP, and none of them is predictive.

Here we present a method, SeqDYN, for predicting *R*_2_ of IDPs. Using a mathematical model introduced by Li et al. (18) to predict propensities for binding nanoparticles and also adapted for predicting propensities for binding membranes (19), we express the *R*_2_ value of a residue as the product of contributing factors from all residues. The contributing factor attenuates as the neighboring residue becomes more distant from the central residue. The model, after training on a set of 45 IDPs, has prediction accuracy that is competitive against that of the recent MD simulations using IDP-specific force fields (1, 12-14). For lysozyme, the SeqDYN prediction agrees remarkably well with *R*_2_ measured in the unfolded state.

## Results

### The data set of IDPs with *R*_2_ rates

We collected *R*_2_ data for a total of 54 nonhomologous IDPs or IDRs (Table 1; Figure 1). According to indicators from NMR properties, including low or negative NOEs, narrow dispersion in backbone amide proton chemical shifts, and small secondary chemical shifts (SCSs), most of the proteins are disordered with at most transient α-helices. A few are partially folded, including Sev-NT with a well-populated (∼80%) long helix (residues 478-491) (20), CREB-binding protein fourth intrinsically disordered linker (CBP-ID4) with > 50% propensities for two long helices (residues 2-25 and 101-128) (21), HOX transcription factor DFD (HOX-DFD) with a well-folded domain comprising three helices (22), and Hahellin (apo form) as a molten globule (23). In Figure 2, we display representative conformations of five IDPs, ranging from fully disordered MAPK kinase 4 (MKK4) (24) and α-synuclein (25) to Measles virus phosphoprotein N-terminal domain (Mev-P_NTD_) (26) with transient short helices to Sev-NT and CBP-ID4 with stable long helices. The sequences of all the IDPs are listed in Table 2.

**Figure 1.**
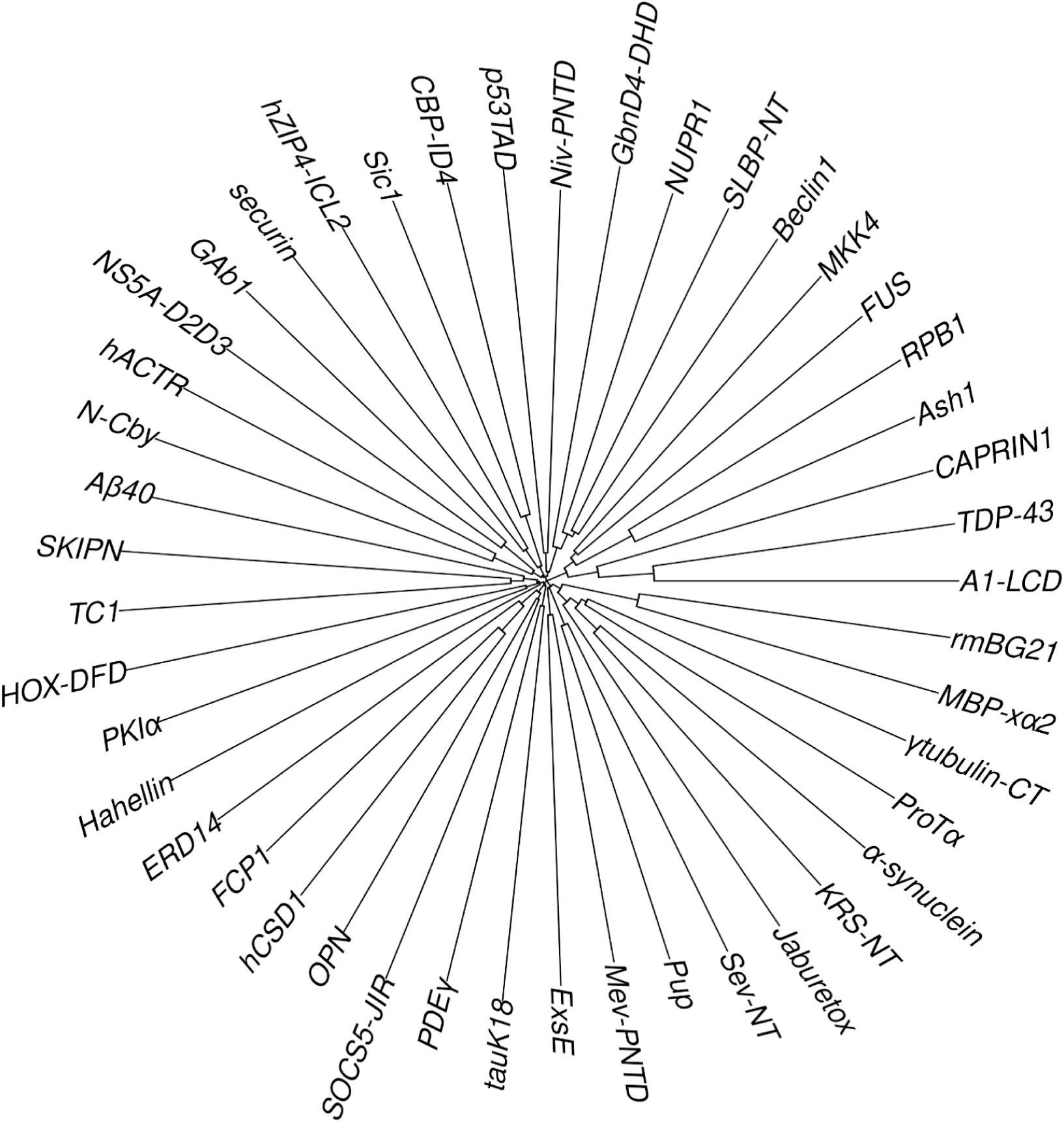
Clock-like tree plot showing lack of homology among the 45 IDPs. The level of homology between two sequences is measured by the distance from their convergence point to the center of the clock. The highest level of apparent identity is between A1-LCD and TDP-43, at 25%, but these two proteins differ in both secondary structure formation and *R*_2_ characteristics. There is, however, a 20-residue overlap between the N-terminus of MBP-xα2 and the C-terminus of rmBG21.

**Figure 2.**
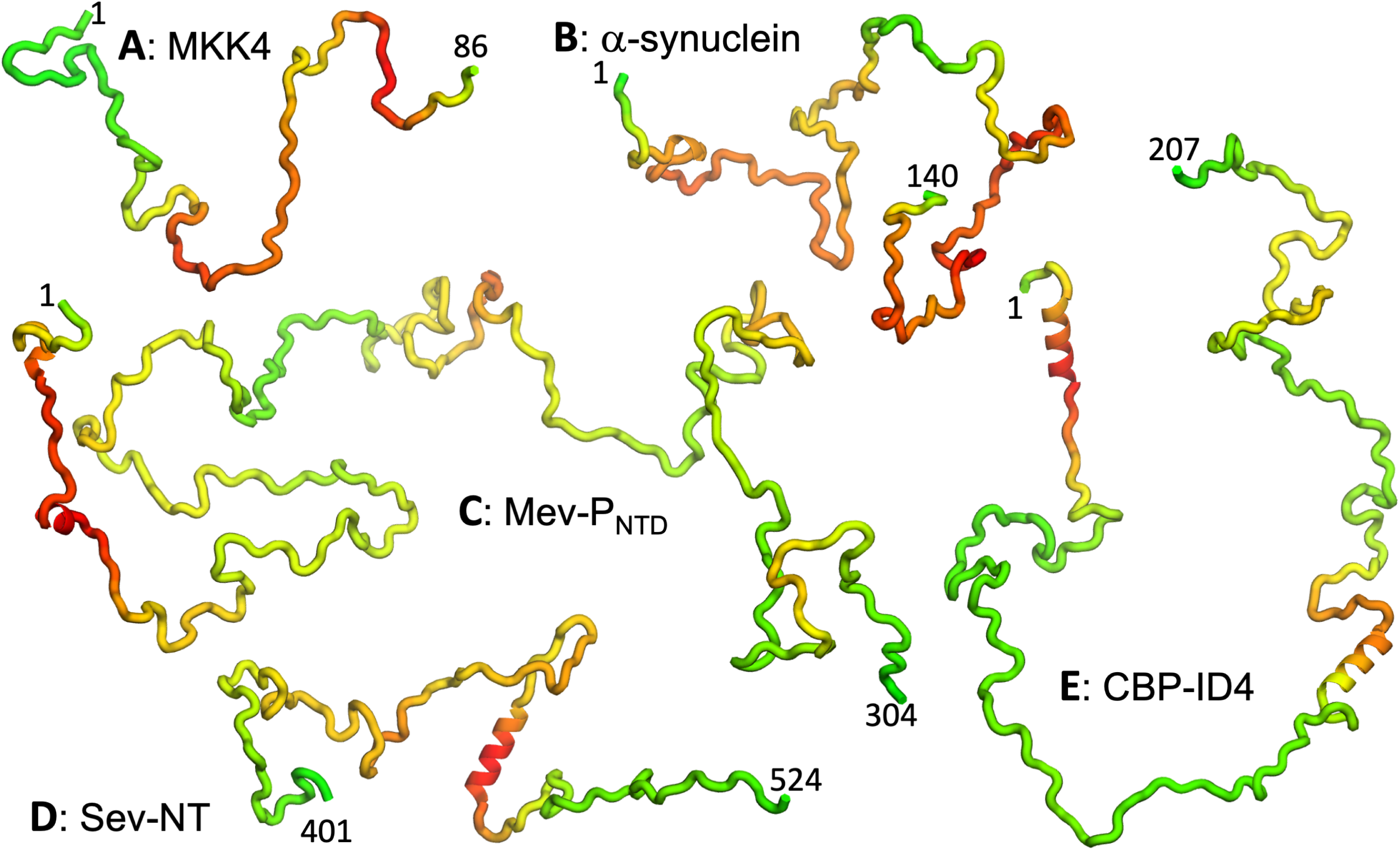
Representative conformations of five IDPs. (A-E) MKK4, α-synuclein, Mev-P_NTD_, Sev-NT, and CBP-ID4. Conformations were initially generated using TraDES (http://trades.blueprint.org) (81), selected to have radius of gyration close to predicted by a scaling function *R*_g_ = 2.54*N*^0.522^ (Å) (82). Conformations for residues predicted as helical by PsiPred plus filtering were replaced by an ideal helix. Finally residues are colored according to a scheme ranging from green for low predicted *R*_2_ to red for high predicted *R*_2_.

**Table 1.**
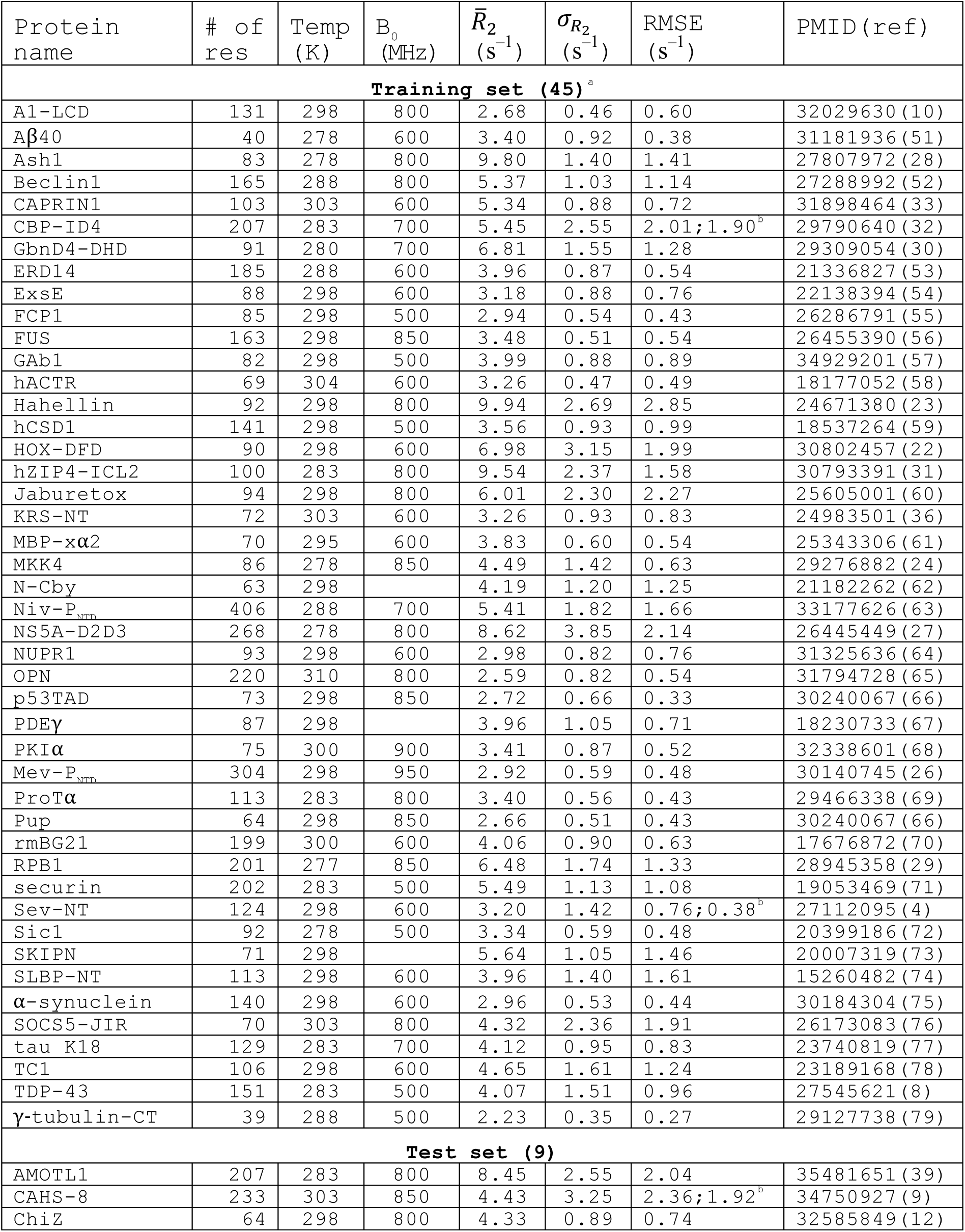
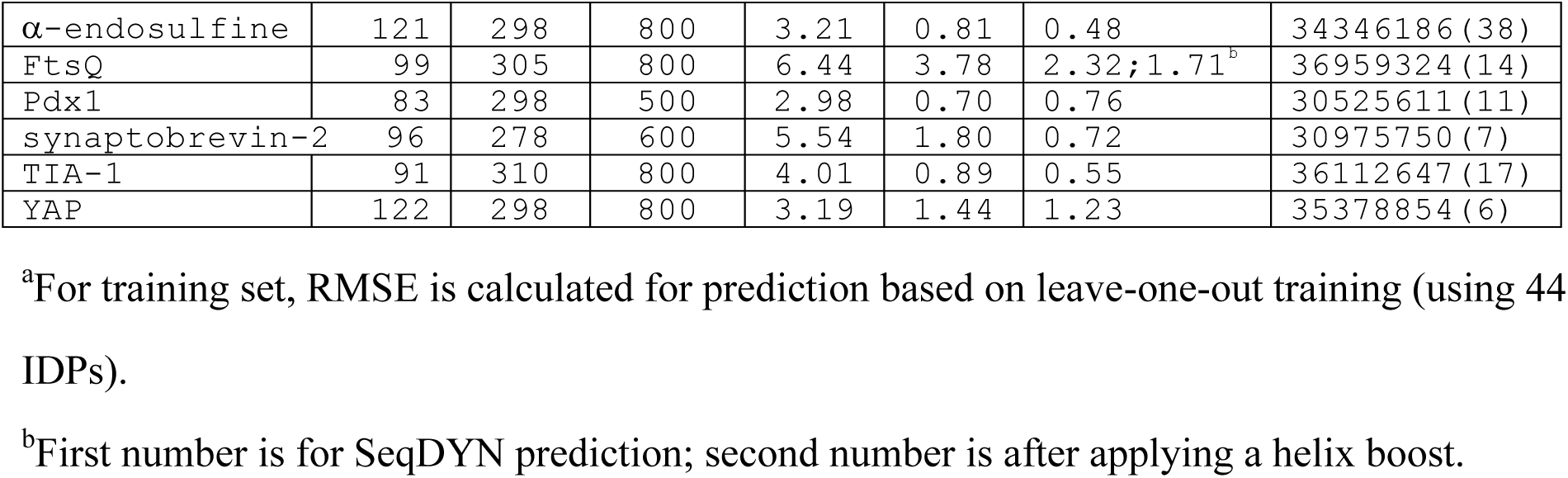
Experimental conditions, means and standard deviations of measured *R*_2_, and SeqDYN prediction RMSEs.

**Table 2.**
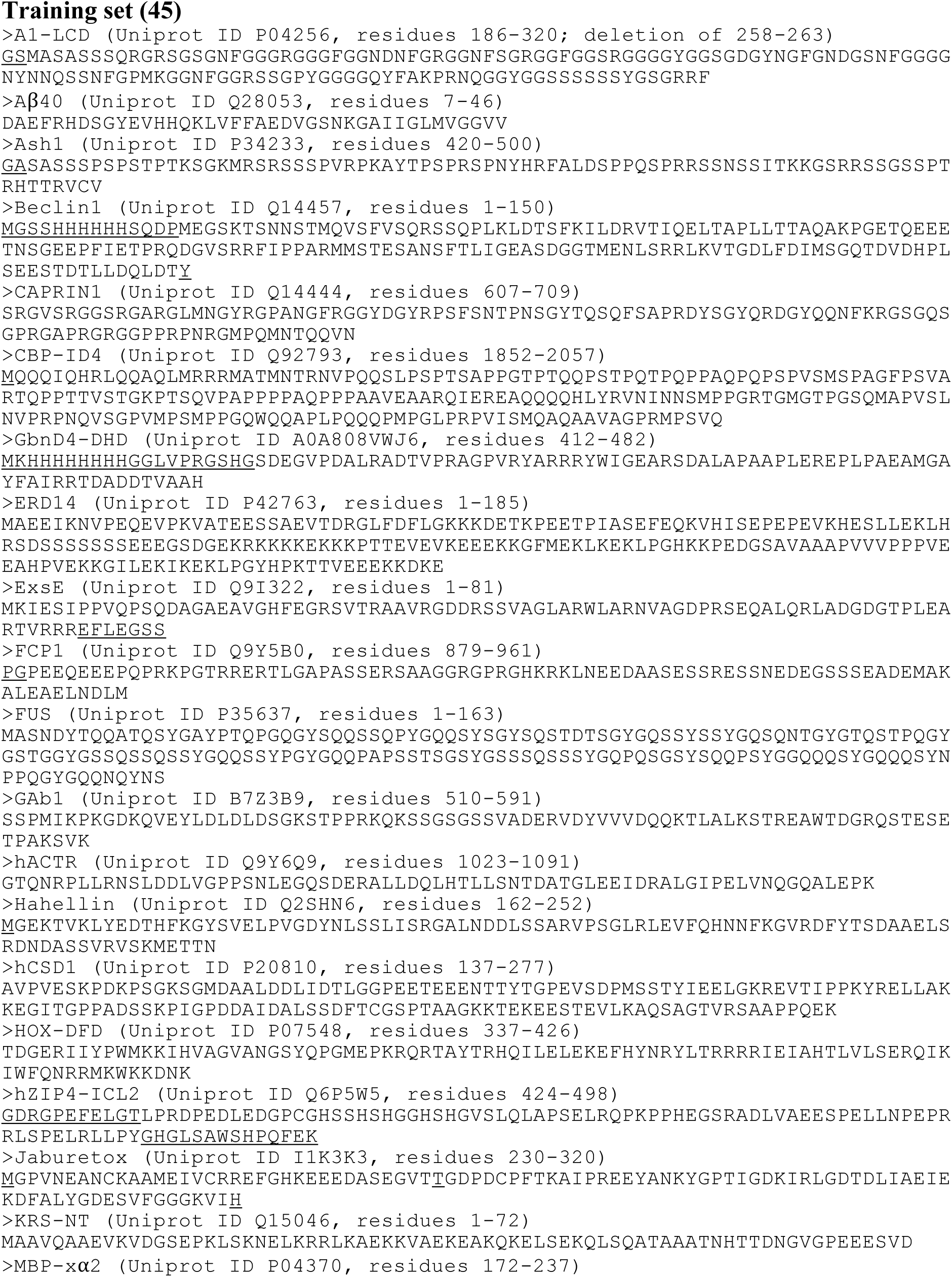
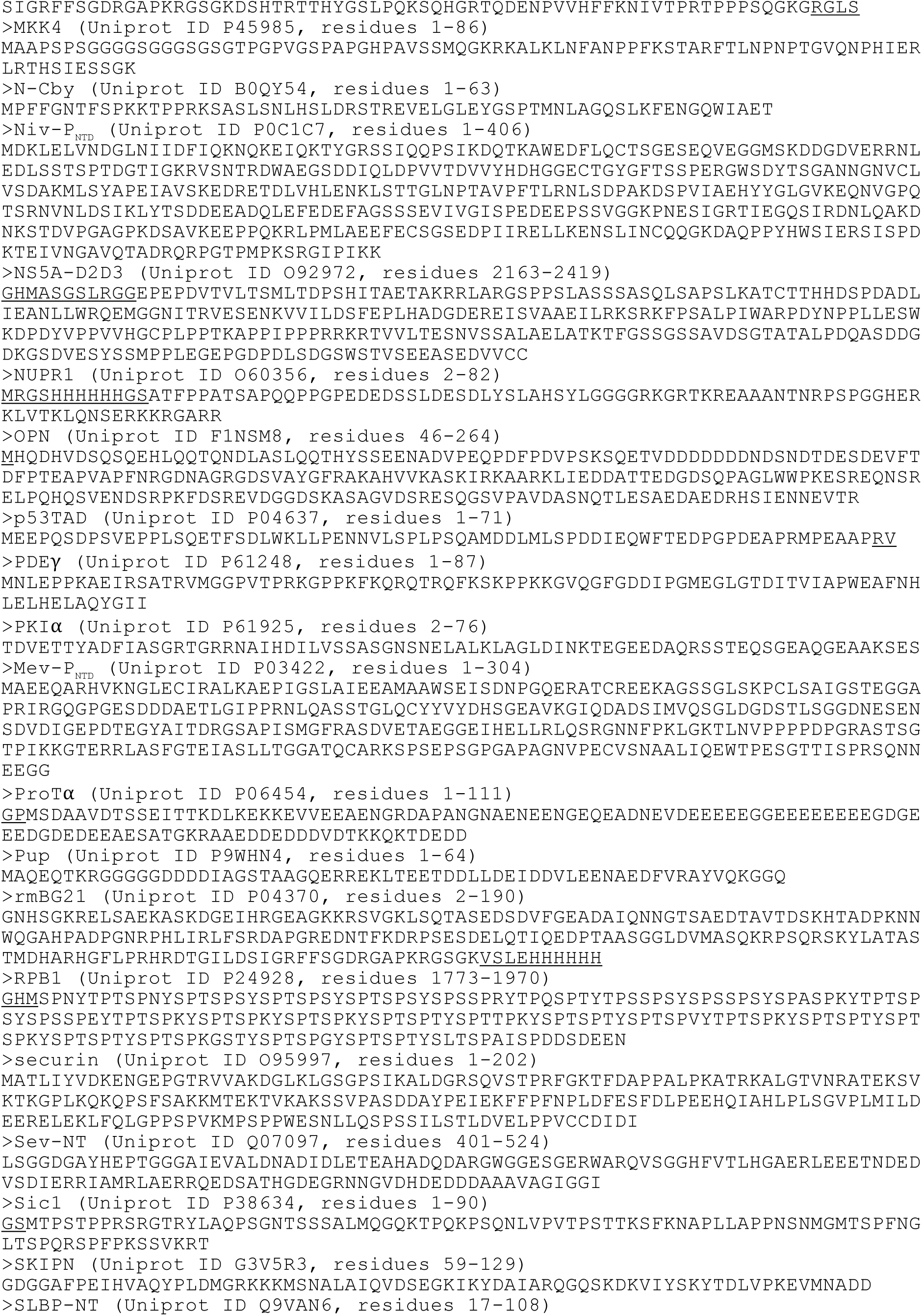
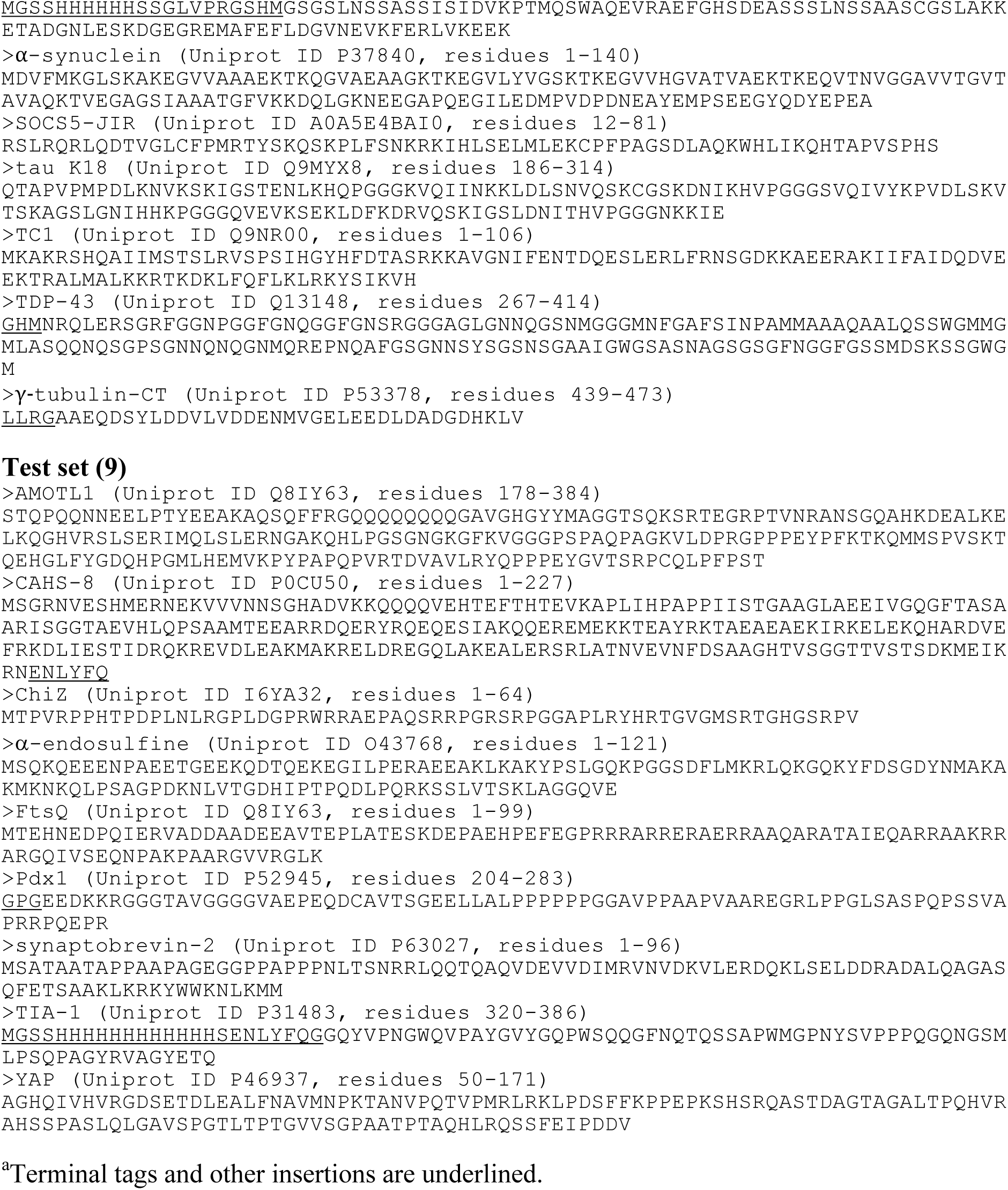
Sequences of 54 IDPs^a^.

We used 45 of the 54 IDPs to train and validate SeqDYN and reserved the remaining 9 for testing. The sequence lengths of the training set range from 39 to 406 residues, with an average of 125.3 residues. Altogether *R*_2_ data are available for 3966 residues. A large majority (35 out of 45) of the 45 IDPs have mean *R*_2_ values (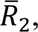 calculated among all the residues in a protein) between 2.5 and 5.5 s^-1^ (Table 1 and Figure 3A). This 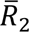 range is much lower than that of structured proteins with similar sequence lengths. The low 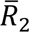 values and lack of dependence on sequence length (Figure 3-figure supplement 1A) suggest that *R*_2_ of the IDPs is mostly dictated by local sequence instead of tertiary interaction.

**Figure 3.**
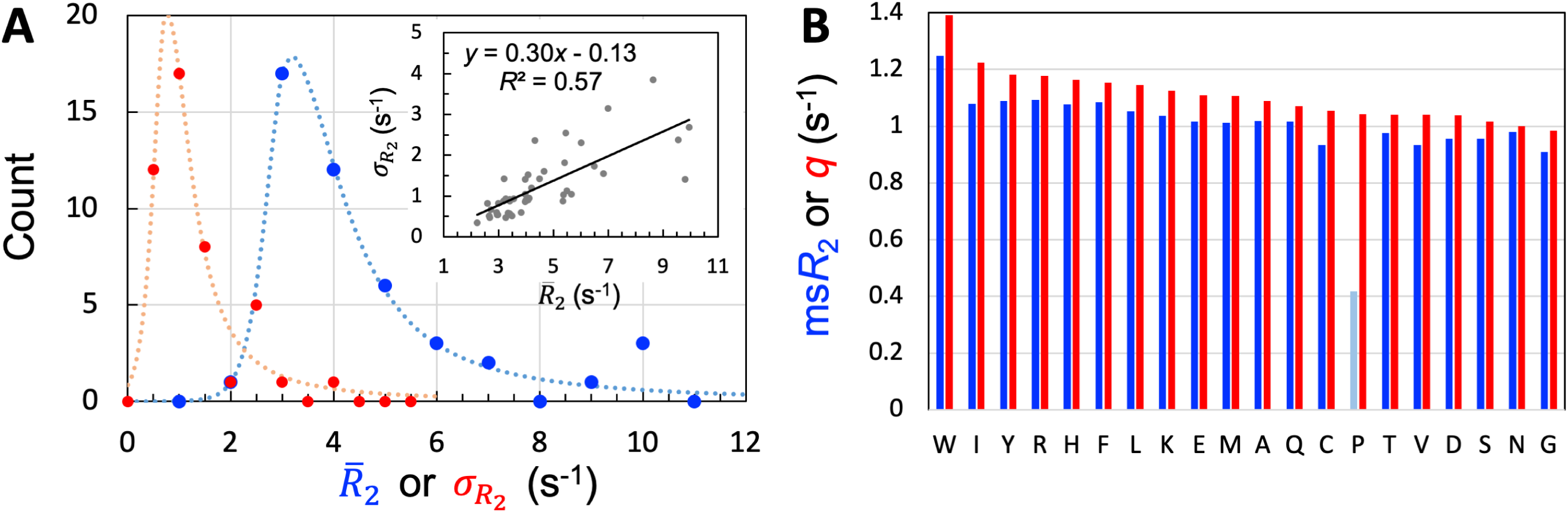
Properties of the 45 IDPs in the training set. (A) Histograms of means and standard deviations, calculated for individual proteins. Curves are drawn to guide the eye. Inset: correlation between 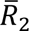 and *σ*_*R*_2__. (B) Experimental mean scaled *R*_2_ (ms*R*_2_) and SeqDYN *q* parameters, for the 20 types of amino acids. Note that Pro residues have low ms*R*_2_ for the lack of backbone amide proton. Amino acids are in descending order of *q*.

The most often used temperature for acquiring the *R*_2_ data was 298 K, but low temperatures (277 to 280 K) were used in a few cases (Table 1 and Figure 3-figure supplement 1B). Of the seven IDPs with 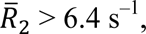 four can be attributed to low temperatures (27-30), one is due to a relatively low temperature (283 K) as well as the presence of glycerol (20% v/v) (31), and two can be explained by tertiary structure formation [a folded domain (22) or molten globule (23)]. A simple reason for higher *R*_2_ values at lower temperatures is the higher water viscosity, resulting in a slowdown in molecular tumbling; a similar effect is achieved by adding glycerol. In some cases, *R*_2_ was measured at both low and room temperatures (4, 10). To a good approximation, the effect of lowering temperature is a uniform scaling of *R*_2_ across the IDP sequence. For Sev-NT, downscaling of the *R*_2_ values at 278 K by a factor of 2.0 brings them into close agreement with those at 298 K (Figure 3-figure supplement 1C), with a root-mean-square-deviation (RMSD) of 0.5 s^-1^ among all the residues. Likewise, for A1-LCD, downscaling by a factor of 2.4 brings the *R*_2_ values at 288 K into good match with those at 298 K (Figure 3-figure supplement 1D), with an RMSD of 0.4 s^-1^. Because SeqDYN is concerned with the sequence dependence of *R*_2_, a uniform scaling has no effect on model parameter or prediction; therefore mixing the data from different temperatures is justified. The same can be said about the different magnetic fields in acquiring the *R*_2_ data (Table 1 and Figure 3-figure supplement 1E). Increasing the magnetic field raises *R*_2_ values, and the effect is also approximated well by a uniform scaling (4, 8, 29).

One measure on the level of sequence dependence of *R*_2_ is the standard deviation, *σ*_*R*_2__, calculated among the residues of an IDP. Among the training set, the *R*_2_ values of 30 IDPs have moderate sequence variations, with *σ*_*R*_2__ ranging from 0.5 to 1.5 s^-1^ (Table 1); the histogram of *σ*_*R*_2__ calculated for the entire training set peaks around 0.75 s^-1^ (Figure 3A). There is a moderate correlation between *σ*_*R*_2__ and 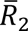 (Figure 3A, inset), reflecting in part the fact that *σ*_*R*_2__ can be raised simply by a uniform upscaling, e.g., as a result of lowering temperature. Still, only two of the five IDPs with high 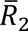 attributable to lower temperature or presence of glycerol are among the seven IDPs with high sequence variations (*σ*_*R*_ > 2 s^-1^). Therefore the sequence variation of *R*_2_ as captured by *σ*_*R*_2__ manifests mostly the intrinsic effect of the IDP sequence, not the influence of external factors such as temperature or magnetic field strength. The mean *σ*_*R*_ value among the training set is 1.24 s^-1^.

One way to eliminate the influence of external factors is to scale the *R*_2_ values of each IDP by its 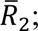 we refer to the results as scaled *R*_2_, or s*R*_2_. We then pooled the s*R*_2_ values for all residues in the training set, and separated them according to amino-acid types. The amino acid type-specific mean s*R*_2_ values, or ms*R*_2_, are displayed in Figure 3B. The seven amino acids with the highest ms*R*_2_ in descending order are Trp, Arg, Tyr, Phe, Ile, His, and Leu. The presence of all the four aromatic amino acids in this “high-end” group immediately suggests π-π stacking as important for raising *R*_2_; the presence of Arg further implicates cation-π interactions. In the other extreme, the seven amino acids with the lowest ms*R*_2_ in ascending order are Gly, Cys, Val, Asp, Ser, Thr, and Asn. Gly is well-known as a flexible residue; it is also interesting that all the four amino acids with short polar sidechains are found in this “low-end” group. Pro has an excessively low ms*R*_2_ [with data from only two IDPs (32, 33)], but that is due to the absence of an amide proton.

### The SeqDYN model and parameters

The null model is to assume a uniform *R*_2_ for all the residues in an IDP. The root-mean-square-error (RMSE) of the null model is equal to the standard deviation, *σ*_*R*_2__, of the measured *R*_2_ values. The mean RMSE, 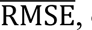 of the null model, equal to 1.24 s^-1^ for the training set, serves as the upper bound for evaluating the errors of *R*_2_ predictors. The next improvement is a one-residue predictor, where first each residue (with index *n*) assumes its amino acid-specific mean s*R*_2_ (ms*R*_2_) and then a uniform scaling factor Υ is applied:

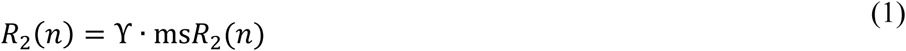

This one-residue model does only minutely better than the null model, with a 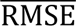 of 1.22 s^-1^.

In SeqDYN, we account for the influence of neighboring residues. Specifically, each residue *i* contributes a factor *f*(*i*; *n*) to the *R*_2_ value of residue *n*. Therefore

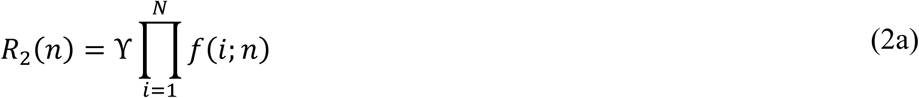

where *N* is the total number of residues in the IDP. The contributing factor depends on the sequence distance *s* = |*i* − *n*| and the amino-acid type of residue *i*:

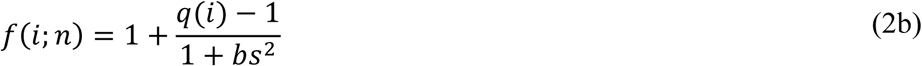

There are 21 global parameters. The first 20 are the *q* values, one for each of the 20 types of amino acids; the last parameter is *b*, appearing in the Lorentzian form of the sequence-distance dependence. We define the correlation length, *L*_corr_, as the sequence distance at which the contributing factor is midway between the values at *s* = 0 and ∞. It is easy to verify that *L*_corr_ = *b*^−1/2^. Note that the single-residue model can be seen as a special case of SeqDYN, with *L*_corr_ set to 0 and *q* set to ms*R*_2_.

The functional forms of Eqs (2a,b) were adapted from Li et al. (18); we also used them for predicting residue-specific membrane association propensities of IDPs (19). In these previous applications, a linear term was also present in the denominator of Eq (2b). In our initial training of SeqDYN, the coefficient of the linear term always converged to near zero. We thus eliminated the linear term. In addition to the Lorentzian form, we also tested a Gaussian form for the sequence-distance dependence and found somewhat worse performance. The more gradual attenuation of the Lorentzian form with increasing sequence distance evidently provides an overall better model for the *R*_2_ data in the entire training set. Others (16, 17, 24) have modeled *R*_2_ as the average of some parameters over a window; a window has an extremely abrupt sequence-distance dependence (1 for *s* < *L*_corr_ and 0 for *s* > *L*_corr_).

We parametrized the SeqDYN model represented by Eqs (2a,b) on the training set of 45 IDPs. In addition to the 21 global parameters noted above, there are also 45 local parameters, namely one uniform scaling factor (Υ) per IDP. The parameter values were selected to minimize the sum of the mean-square-errors for the IDPs in the training set, calculated on *R*_2_ data for a total of 3924 residues. We excluded the 42 Pro residues in the training set because, as already noted, their *R*_2_ values are lower for chemical reasons. We will present validation and test results below, but first let us look at the parameter values.

The *q* values are displayed in Figure 3B alongside ms*R*_2_. In descending order, the seven amino acids with the highest *q* values are Trp, Ile, Tyr, Arg, His, Phe, and Leu. These are exactly the same amino acids in the high-end group for ms*R*_2_, though their order there is somewhat different. In ascending order, the seven amino acids (excluding Pro) with the lowest *q* values are Gly, Asn, Ser, Asp, Val, Thr, and Cys. The composition of the low-end group is also identical to that for ms*R*_2_. The *q* values thus also suggest that π-π and cation-π interactions in local sequences may raise *R*_2_, whereas Gly and short-polar residues may lower *R*_2_.

Given the common amino acids at both the high and low ends for ms*R*_2_ and *q*, it is not surprising that these two properties exhibit a strong correlation, with a coefficient of determination (*R*^2^; excluding Pro) at 0.92 (Figure 4A). Also, because the high-end group contains the largest amino acids (e.g., Trp and Tyr) whereas the low-end group contains the smallest amino acids (e.g., Gly and Ser), we anticipated some correlation of ms*R*_2_ and *q* with amino-acid size. We measure the latter property by the molecular mass (*m*). As shown in Figure 4B, both ms*R*_2_ and *q* indeed show a medium correlation with *m*, with *R*^2^ = 0.67 (excluding Pro) and 0.61, respectively. A bulkiness parameter was proposed as an indicator of sequence-dependent backbone dynamics of IDPs (16, 24). Bulkiness was defined as the sidechain volume-to-length ratio, and identified amino acids with aromatic or branched aliphatic sidechains as bulky (34). We found only modest correlations between either ms*R*_2_ or *q* and bulkiness, with *R*^2^ just below 0.4 (Figure 4C).

**Figure 4.**
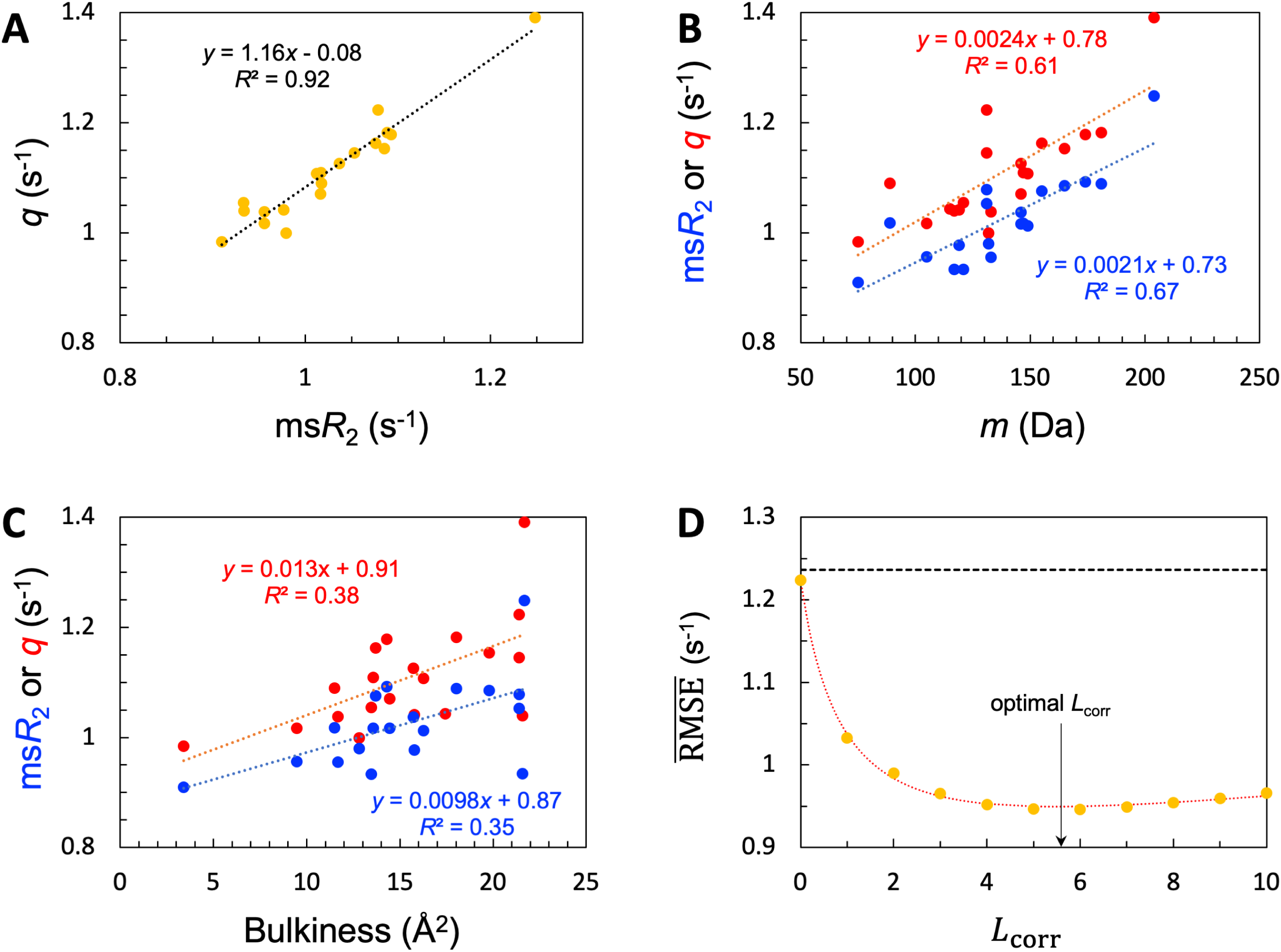
SeqDYN model parameters. (A) Correlation between ms*R*_2_ and *q*. The values are also displayed as bars in Figure 3B. (B) Correlation of ms*R*_2_ and *q* with amino-acid molecular mass. (C) Correlation of ms*R*_2_ and *q* with bulkiness. (D) The optimal correlation length and deterioration of SeqDYN prediction as the correlation length is moved away from the optimal value.

The optimized value of *b* is 3.164 × 10^-2^, corresponding to an *L*_corr_ of 5.6 residues. The resulting optimized 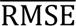 is 0.95 s^-1^, a clear improvement over the value 1.24 s^-1^ of the null model. To check the sensitivity of prediction accuracy to *b*, we set *b* to values corresponding to *L*_corr_ = 0, 1, 2, .., and retrained SeqDYN for *b* fixed at each value. Note that the null-model 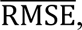 1.24 s^-1^, sets an upper bound. This upper bound is gradually reached when *L*_corr_ is increased from the optimal value. In the opposite direction, when *L*_corr_ is decreased from the optimal value, 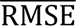 rises quickly, reaching 1.22 s^-1^ at *L*_corr_ = 0. The latter 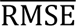 is the same as that of the single-residue model. Lastly we note that there is a strong correlation between the uniform scaling factors and 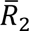 values among the 45 IDPs (*R*^2^ = 0.77), as to be expected. For 39 of the 45 IDPs, Υ values fall in the range of 0.8 to 2.0 s^-1^.

As presented next, we evaluate the performance of SeqDYN by leave-one-out cross validation, where each IDP in turn was left out of the training set and the model was trained on the remaining 44 IDPs to predict *R*_2_ for the IDP that was left out. The parameters from the leave-one-out (also known as jackknife) training sessions allow us to assess the potential bias of the training set. For this purpose, we compare the values of the 21 global parameters, either from the full training set or from taking the averages of the jackknife training sessions. For each of the *q* parameters, the values from these two methods differ only in the fourth digit; e.g., for Leu, they are both 1.1447 from full training and from jackknife training. The values for *b* are 3.164 × 10^-2^ from full training as stated above and 3.163 × 10^-2^ from jackknife training. The close agreement in parameter values between full training and jackknife training suggests no significant bias in the training set.

Another question of interest is whether the difference between the *q* parameters of two amino acids is statistically significant. To answer this question, we carried out five-fold cross-validation training, resulting in five independent estimates for each parameter. For example, the mean ± standard deviation of the *q* parameter is 1.1405 ± 0.0066 for Leu and 1.2174 ± 0.0211 for Ile. A t-test shows that their difference is extremely statistically significant (*P* < 0.0001). In contrast, the difference between Leu and Phe (*q* = 1.1552 ± 0.0304) is not significant. t-test results for other pairs of amino acids are found in Figure 4-figure supplement 1.

### Validation of SeqDYN predictions

We now present leave-one-out cross-validation results. We denote the RMSE of the *R*_2_ prediction for the left-out IDP as RMSE(-1). As expected, RMSE(-1) is higher than the RMSE obtained with the IDP kept in the training set, but the increases are generally slight. Specifically, all but eight of the IDPs have increases < 0.1 s^-1^; the largest increase is 0.35 s^-1^, for CBP-ID4. The mean RMSE(-1), or 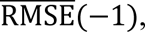 for the 45 IDPs is increased by 0.05 s^-1^ over 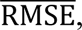 to 1.00 s^-1^. The latter value is still a distinct improvement over the mean RMSE 1.24 s^-1^ of the null model. The histogram of RMSE(-1) for the 45 IDPs is shown in Figure 5A. It peaks at 0.5 s^-1^, which is a substantial downshift from the corresponding peak at 0.75 s^-1^ for *σ*_*R*_2__ (Figure 3A). Thirty-four of the 45 IDPs have RMSE(-1) values lower than the corresponding *σ*_*R*_2__.

**Figure 5.**
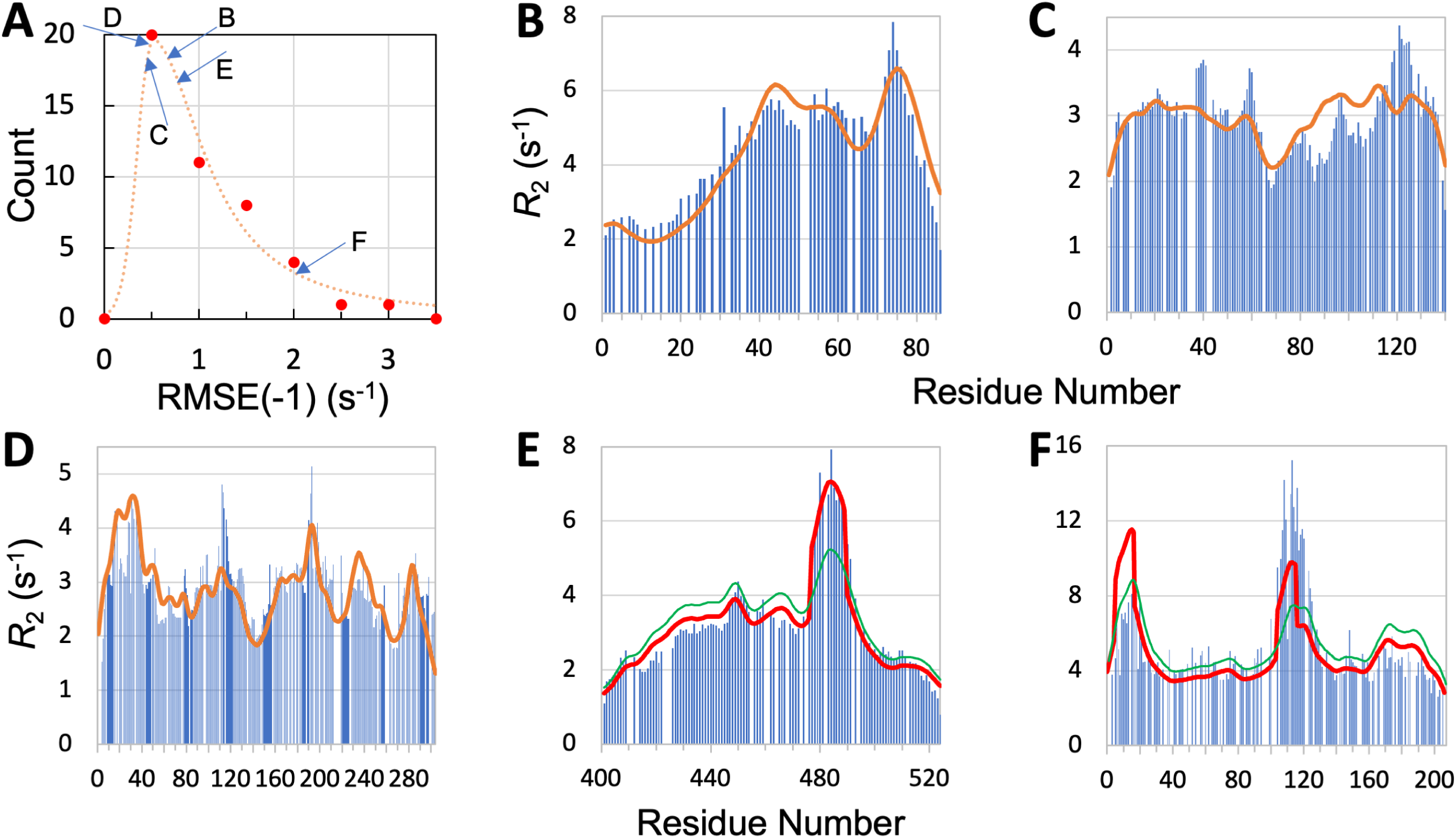
Quality of SeqDYN predictions. (A) Histogram of RMSE(-1). Letters indicate RMSE(-1) values of the IDPs to be presented in panels (B-F). (B-F) Measured (bars) and predicted (curves) *R*_2_ profiles for MKK4, α-synuclein, Mev-P_NTD_, Sev-NT, and CBP-ID4. In (E) and (F), green curves are SeqDYN predictions and red curves are obtained after a helix boost.

To further illustrate the performance of SeqDYN, we present the comparison of predicted and measured *R*_2_ values for five IDPs: MKK4, α-synuclein, Mev-P_NTD_, Sev-NT, and CBP-ID4 (Figure 5B-F). A simple common feature is the falloff of *R*_2_ at the N- and C-termini, resulting from missing upstream or downstream residues that otherwise would be coupled to the terminal residues, as first recognized by Schwalbe et al. (15). Representative conformations of the five IDPs are displayed in Figure 2, with residues colored according to the predicted *R*_2_ values. For four of these IDPs, the RMSE(-1) values range from 0.44 to 0.76 s^-1^ and are scattered around the peak of the histogram, while the RMSE(-1) for the fifth IDP, namely CBP-ID4, the RMSE(-1) value is 2.01 s^-1^ and falls on the tail of the histogram (Figure 5A). Figure 5B displays the measured and predicted *R*_2_ for MKK4. SeqDYN correctly predicts higher *R*_2_ values in the second half of the sequence than in the first half. It even correctly predicts the peak around residue Arg75. The sequence in this region is H_72_IERLRTH_79_; six of these eight residues belong to the high-end group. In contrast, the lowest *R*_2_ values occur in the sequence S_7_GGGGSGGGSGSG_19_, comprising entirely of two amino acids in the low-end group.

*R*_2_ values for α-synuclein are shown in Figure 5C. Here SeqDYN correctly predicts higher *R*_2_ near the C-terminus and a dip around Gly68. However, it misses the *R*_2_ peaks around Tyr39 and Asp121. MD simulations (1) have found that these *R*_2_ peaks can be explained by a combination of secondary structure formation (β-sheet around Tyr39 and polyproline II helix around Asp121) and local (between Tyr39 and Ser42) or long-range (between Asp121 and Lys96) interactions. SeqDYN cannot account for long-range interactions (e.g., between β-strands and between Asp121 and Lys96). Figure 5D shows that SeqDYN gives excellent *R*_2_ predictions for Mev-P_NTD_. It correctly predicts the high peaks around Arg17, Glu31, Leu193, and lower peaks around Arg235 and Trp285, but does underpredict the narrow peak around Tyr113.

The overall *R*_2_ profile of Sev-NT is predicted well by SeqDYN, but the peak in the long helical region (residues 478-491) is severely underestimated (green curve in Figure 5E). A similar situation occurs for CBP-ID4, where the peak in the second long helical region (around Glu113) is underpredicted (green curve in Figure 5F). While the measured *R*_2_ exhibits a higher peak in the second helical region than in the first helical region (around Arg16), the opposite is predicted by SeqDYN. When the *R*_2_ data were included in the training set (i.e., full training), the second peak is higher than the first one, but that is not a real prediction because the *R*_2_ data themselves were used for training the model. It merely means that the SeqDYN functions can be parameterized to produce any prescribed *R*_2_ profile along the sequence. Indeed, when the *R*_2_ data of CBP-ID4 alone were used to parameterize SeqDYN, the measured *R*_2_ profile is closely reproduced (Figure 5-figure supplement 1). The reversal in *R*_2_ peak heights between the two helical regions is the reason for the aforementioned unusual increase in RMSE when CBP-ID4 was left out of the training set.

### R2 boost in long helical regions

It is apparent that SeqDYN underestimates the *R*_2_ of stable long helices. Transient short helices does not seem to be a problem, since these are present, e.g., in Mev-P_NTD_, where transient helix formation in the first 37 residues and between residues 189-198 (26) coincides with *R*_2_ peaks that are correctly predicted by SeqDYN. SeqDYN can treat coupling between residues within the correlation length of 5.6 residues, but a much longer helix would tumble more slowly than implied by an *L*_corr_ of 5.6, and thus it makes sense that SeqDYN would underestimate *R*_2_ in that case.

Our solution then is to apply a boost factor to the long helical region. To do so, we have to know whether an IDP does form long helices and if so what the constituent residues are. Secondary structure predictors tend to overpredict α-helices and β-strands for IDPs, as they are trained on structured proteins. One way to counter that tendency is to make the criteria for α-helices and β-strands stricter. We found that, by filtering PsiPred (http://bioinf.cs.ucl.ac.uk/psipred) (35) helix propensity scores (*p*_Hlx_) with a very high cutoff of 0.99, the surviving helix predictions usually correspond well with residues identified by NMR as having high helix propensities. For example, for Mev-P_NTD_, PsiPred plus filtering predicts residues 14-17, 28-33, and 191-193 as helical; all of them are in regions that form transient helices according to chemical shifts (26). Likewise long helices are also correctly predicted for Sev-NT (residues 477-489) and CBP-ID4 (residues 6-17 and 105-116) (20, 21).

We apply a boost factor, *B*_Hlx_, to helices with a threshold length of 12:

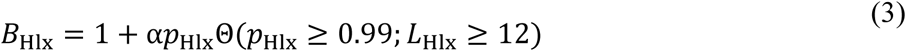

The Θ function is 1 if the helix propensity score is above the filtering cutoff and the helix length (*L*_Hlx_) is above the threshold, and 0 otherwise. With a boost amplitude α at 0.5, the boosted SeqDYN prediction for Sev-NT reaches excellent agreement with the measured *R*_2_(Figure 5E, red curve). The RMSE(-1) is reduced from 0.76 s^-1^ to 0.38 s^-1^ upon boosting. Applying the same helix boost to CBP-ID4 also results in a modest reduction in RMSE(-1), from 2.01 to 1.90 s^-1^ (Figure 5F, red curve). The only other IDP for which PsiPred plus filtering predicts a long helix is the N-terminal region of lysyl-tRNA synthetase (KRS-NT). The authors who studied this protein did not report on secondary structure (36), but feeding their reported chemical shifts to the TALOS+ server (https://spin.niddk.nih.gov/bax/nmrserver/talos/) (37) found only short stretches of residues that fall into the helical region of the Ramachandran map. The SeqDYN prediction for KRS-NT is already good [RMSE(-1) = 0.83 s^-1^]; applying a helix boost would deteriorate the RMSE(-1) to 1.16 s^-1^.

### Further test on a set of nine IDPs

We have reserved nine IDPs for testing SeqDYN (parameterized on the training set of 45 IDPs). The level of disorder in these test proteins also spans the full range, from absence of secondary structures [ChiZ N-terminal region (12), Pdx1 C-terminal region (11), and TIA-1 prion-like domain (17)] to presence of transient short helices [synaptobrevin-2 (7), α-endosulfine (38), YAP (6), angiomotin-like 1 (AMOTL1) (39)] to formation of stable long helices [FtsQ (14) and CAHS-8 (9)]. For eight of the nine test IDPs, the RMSEs of SeqDYN predictions are lower than the experimental *σ*_*R*_2__ values, by an average of 0.66 s^-1^. For the ninth IDP (Pdx1), the SeqDYN RMSE is slightly higher, by 0.06 s^-1^, than the experimental *σ*_*R*_2__. Together, the nine test IDPs have a mean RMSE of 1.13 s^-1^, close to the 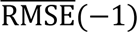 of 1.00 s^-1^ for the training set in the leave-one-out cross-validation.

The comparison of predicted and measured *R*_2_ profiles along the sequence is presented in Figure 6A-I. For ChiZ, SeqDYN correctly predicts the major peak around Arg25 and the minor peak around Arg46 (Figure 6A). The *R*_2_ profile of Pdx1 is largely featureless, except for a dip around Gly216, which is correctly predicted by SeqDYN (Figure 6B). Correct prediction is also obtained for the higher *R*_2_ in the first half of TIA-1 prion-like domain than in the second half (Figure 6C). SeqDYN gives an excellent prediction for synaptobrevin-2, including a linear increase up to Arg56 and the major peak around Trp89 (Figure 6D).

**Figure 6.**
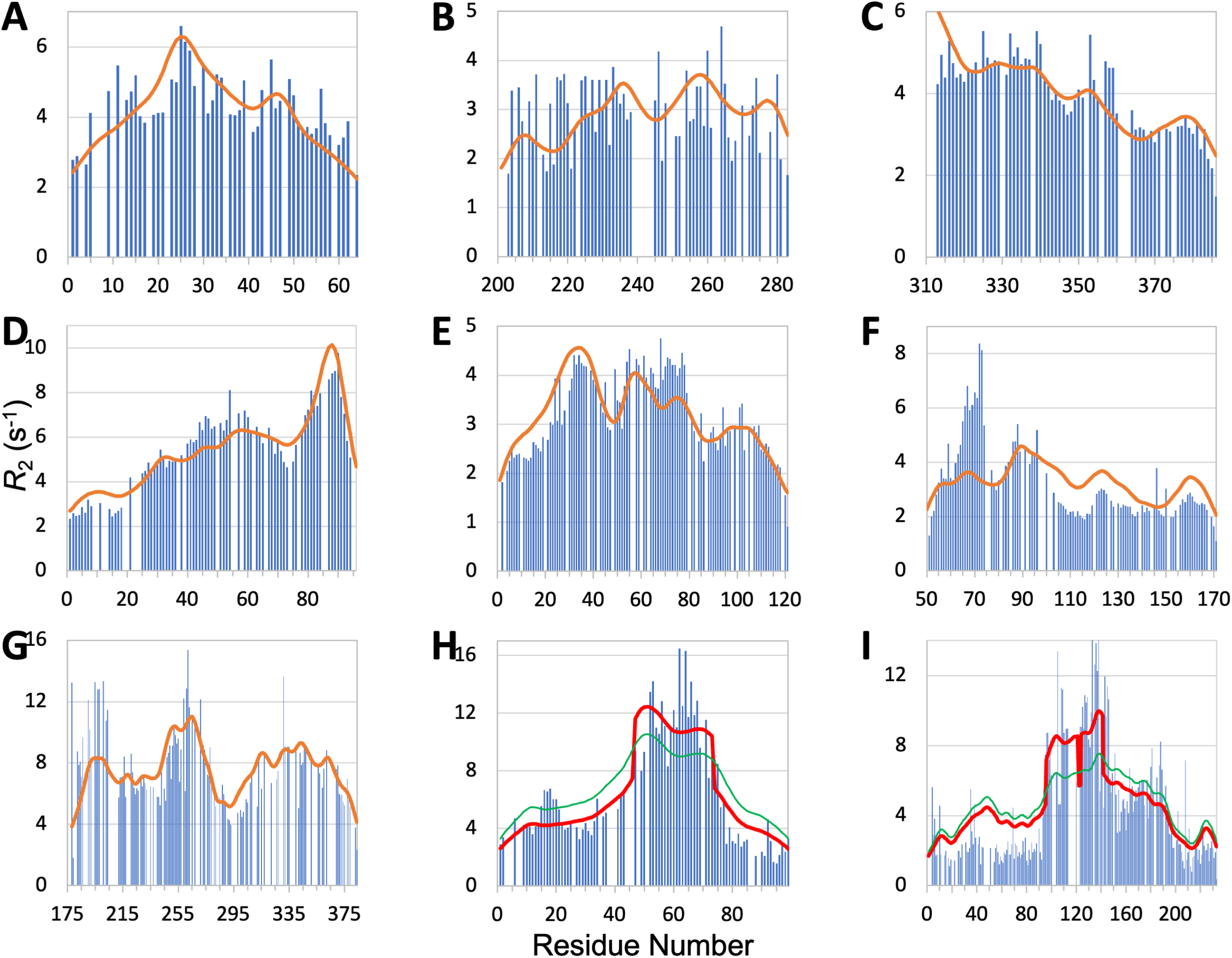
Measured (bars) and predicted (curves) *R*_2_ profiles for ChiZ N-terminal region, TIA1 prion-like domain, Pdx1 C-terminal region, synaptobrevin-2, α-endosulfine, YAP, AMOTL1, FtsQ, and CAHS-8. In (C), *R*_2_ does not fall off at the N-terminus because the sequence is preceded by an expression tag MGSSHHHHHHHHHHHHS. In (H) and (I), green curves are SeqDYN predictions and red curves are obtained after a helix boost.

The prediction is also very good for α-endosulfine, including elevated *R*_2_ around Glu34, which coincides with the presence of a transient helix, and depressed *R*_2_ in the last 40 residues (Figure 6E). The only miss is an underprediction for the peak around Lys74. SeqDYN also predicts well the overall shape of the *R*_2_ profile for YAP, including peaks around Asn70, Leu91, Arg124, and Arg161, but severely underestimates the peak height around Asn70 (Figure 6F).NOE signals indicate contacts between Met86, Leu91, Fhe95, and Fhe96 (6); evidently this type of local contacts is captured well by SeqDYN. The *R*_2_ elevation around Asn70 is mostly due to helix formation: residues 61-74 have helix propensities up to 40% (6). PsiPred predicts helix for residues 62-73, but only residues 65-68 survive the filtering that we impose, resulting in a helix that is too short to apply a helix boost. The prediction for AMOTL1 is mostly satisfactory, including peaks around Phe200 and Arg264 and a significant dip around Gly292 (Figure 6G). However, whereas the two peaks have approximately equal heights in the measured *R*_2_ profile, the predicted peak height around Phe200 is too low. SCSs indicate helix propensity around both *R*_2_ peaks (39). PsiPred also predicts helix in both regions, but only five and two residues, respectively, survive after filtering, and are too short for applying a helix boost.

For FtsQ, SeqDYN correctly predicts elevated *R*_2_ for the long helix [residues 46-74 (14)] but underestimates the magnitude (RMSE = 2.32 s^-1^; green curve in Figure 6H). PsiPred plus filtering predicts a long helix formed by residues 47-73. Applying the helix boost substantially improves the agreement with the measured *R*_2_, with RMSE reducing to 1.71 s^-1^ (red curve in Figure 6H). SeqDYN also gives a qualitatively correct *R*_2_ profile for CAHS-8, with higher *R*_2_ for the middle section (residues 95-190) (RMSE = 2.36 s^-1^; green curve in Figure 6I). However, it misses the extra elevation in *R*_2_ for the first half of the middle section (residues 95-145). According to SCS, the first and second halves have helix propensities of 60% and 30%, respectively (9). PsiPred plus filtering predicts helices for residues 96-121, 124-141, 169-173, and 179-189. Only the first two helices, both in the first half of the middle section, are considered long according to our threshold. Once again, applying the helix boost leads to marked improvement in the predicted in *R*_2_, with RMSE reducing to 1.92 s^-1^ (red curve in Figure 6I).

### Inputting the sequences of structured proteins predicts *R*_2_ in the unfolded state

SeqDYN is trained on IDPs, what if we feed it with the sequence of a structured protein? The prediction using the sequence of hen egg white lysozyme, a well-studied single-domain protein, is displayed in Figure 7A. It shows remarkable agreement with the *R*_2_ profile measured by Klein-Seetharaman et al. (3) in the unfolded state (denatured by 8 M urea at pH 2 and reduced to break disulfide bridges), including a major peak around Trp62, a second peak around Trp111, and a third peak around Trp123. Klein-Seetharaman et al. mutated Trp62 to Gly and the major peak all but disappeared. This result is also precisely predicted by SeqDYN with the mutant sequence (Figure 7B).

**Figure 7.**
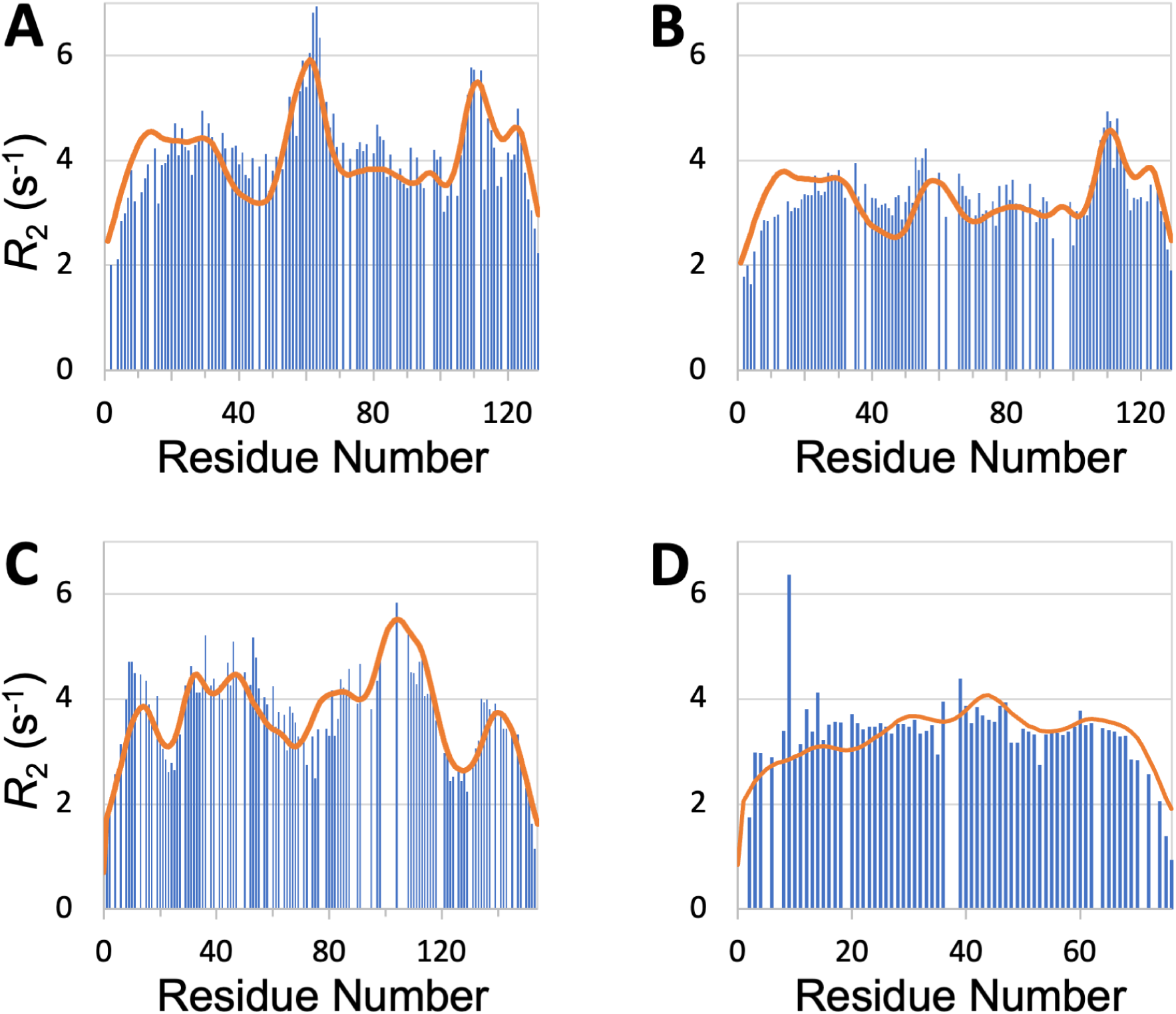
*R*_2_ profiles predicted (curves) by SeqDYN show close agreement with those measured (bars) on structured proteins in the unfolded state. (A) Wild-type lysozyme (8 M urea; pH 2; cysteine-methylated). (B) Lysozyme with Trp62 to Gly mutation (pH 2). Methylated cysteines were treated as Ala in the SeqDYN predictions. (C) Apomyoglogin (8 M urea; pH 2.3). (D) Ubiquitin (8 M urea; pH 2).

SeqDYN also predicts well the *R*_2_ profiles of other proteins in the unfolded state. For unfolded apomyoglobin (8 M urea; pH 2.3), Schwarzinger et al. (40) claimed that depressed *R*_2_ corresponded to stretches of small amino acids (Gly and Ala) whereas elevated corresponded to local hydrophobic interactions. SeqDYN reproduces all the observed peaks and valleys in the *R*_2_ profile (Figure 7C). The deepest valley indeed occurs over a Gly/Ala-rich stretch, <u>G_125_ADAQGA_131_</u>, but the highest peak occurs over a stretch, I_102_KYLEFI_108_, that contains both hydrophobic and charged residues, all of which are on the high end of the *q* parameters (Figure 3B). The *R*_2_ profile of unfolded ubiquitin (8 M urea; pH 2) is relatively flat, which Wirmer et al. (41) attributed to lack of residual secondary structure, based on the assumption that β-sheets (major elements of folded ubiquitin) are less resistant to denaturation than α-helices. SeqDYN predicts a relatively flat *R*_2_ profile (Figure 7D), but the reason is that the ubiquitin sequence lacks a contiguous stretch of high-*q* amino acids.

## Discussion

We have developed a powerful method, SeqDYN, that predicts the backbone amide transverse relaxation rates (*R*_2_) of IDPs. The method is based on IDP sequences, is extremely fast, and available as a web server at https://zhougroup-uic.github.io/SeqDYNidp/. The excellent performance supports the notion that the ns-dynamics reported by *R*_2_ is coded by the local sequence, comprising up to 6 residues on either side of a given residue. The amino-acid types that contribute the most to coupling within a local sequence are aromatic (Trp, Tyr, Phe, and His), Arg, and long branched aliphatic (Ile and Leu), suggesting the importance of π-π, cation-π, and hydrophobic interactions in raising *R*_2_. These interactions are interrupted by Gly and amino acids with short polar sidechains (Ser, Thr, Asn, and Asp), leading to reduced *R*_2_. Transient short helices produce moderate elevation in *R*_2_, whereas stable long helices result in a big boost in *R*_2_. Tertiary contacts can also raise *R*_2_, but appear to be infrequent in most IDPs (1).

It is also possible that *R*_2_ reported by backbone amide ^15^N relaxation (as is the case for most of the IDPs studied here) may not be particularly sensitive to exchange effects, which likely involve tertiary contact formation. For the D2 domain of p27^Kip1^, the exchange contributions measured using ^15^N relaxation were small (< 2.5 s^-1^) but were as large as 25 s^-1^ when measured by high-power ^1^H relaxation dispersion (42). This experiment measures the effective transverse relation rate, *R*_2,eff_, over a range of effective radiofrequency *ω*_eff_. The exchange contribution is maximal for the value 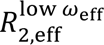 at low *ω*_eff_ but is largely quenched for the value 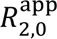 in the high-*ω*_eff_ limit. The SeqDYN prediction for this IDP matches much better with 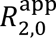 than with 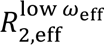 (Figure 8). It is not clear whether this IDP is unique in forming persistent tertiary contacts that give rise to substantial exchange contributions or the ^1^H relaxation dispersion experiment is unique in reporting the exchange contributions. At the minimum, SeqDYN yields the exchange-free portion of the transverse relaxation rate, enabling easy identification of residues that potentially participate in tertiary contacts. For the D2 domain of p27^Kip1^, SeqDYN correctly predicts the 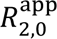 local maxima at W76 and Y88. It is these same two residues that show substantial exchange contributions and putatively participate in tertiary contact (42). Therefore local contacts may seed tertiary contacts. If *R*_2,eff_ data with substantial exchange contributions become available for more IDPs, SeqDYN may be retrained to make predictions for IDPs forming persistent tertiary contacts.

**Figure 8.**
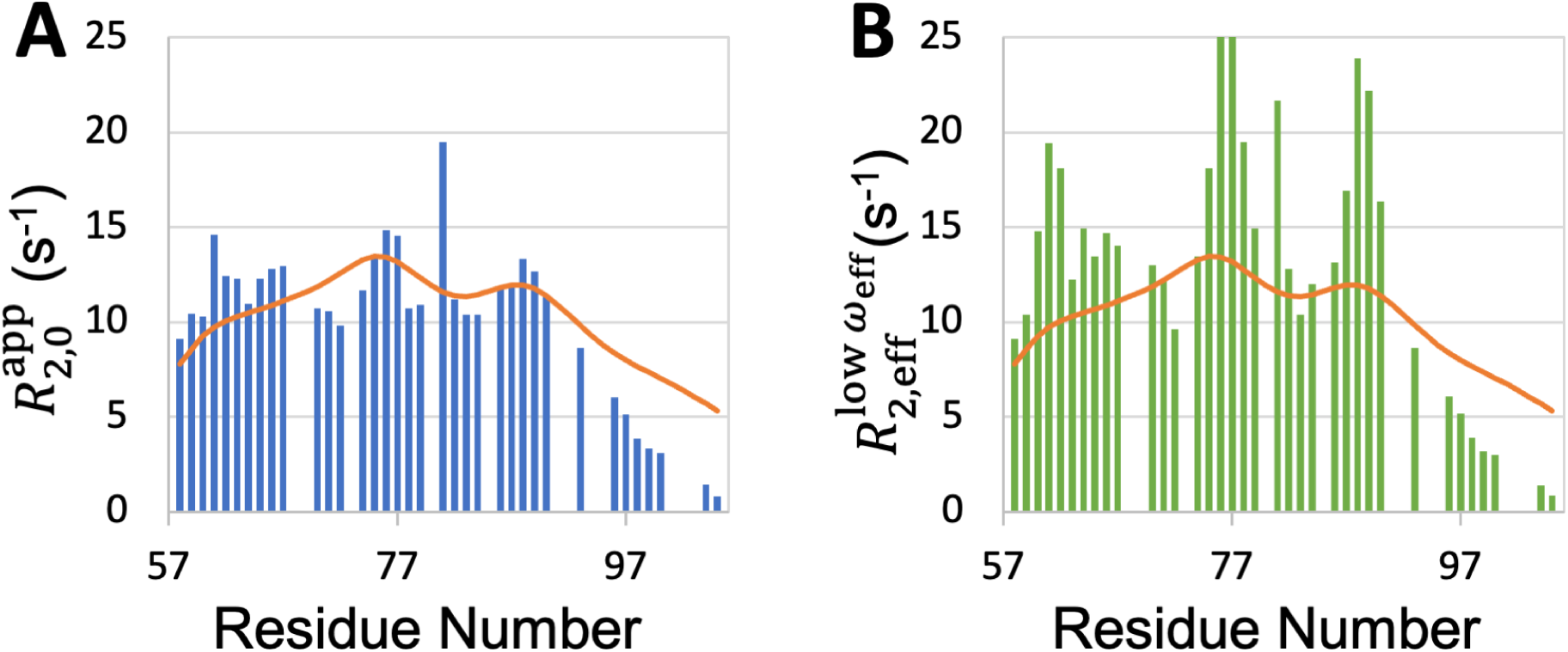
Comparison between SeqDYN prediction (curves) and effective transverse relaxation rate (bars) from ^1^H dispersion relaxation experiment. (A) *R*_2,eff_ in the high-*ω*_eff_ limit. (B) *R*_2,eff_ at low *ω*_eff_.

The *q* parameters, while introduced here to characterize the propensities of amino acids to participate in local interactions, appear to correlate with the tendencies of amino acids to drive liquid-liquid phase separation. Consistent with the rank order of *q*, Trp, Tyr, and Arg have been reported to be strong drivers of phase separation, Lys is a moderate driver, whereas Gly and Ser suppress phase separation (10, 33, 43). Recent measurements of the threshold concentration produced the following order for the propensity of phase separation by eight nonpolar amino acids in homotetrapeptides of the form XXssXX (ss: backbone disulfide bond): Trp > Phe > Leu > Met > Ile > Val > Ala > Pro (44). This order is the same as that of the *q* parameters, except that the *q* values of Ile and Val are in the second and last places, respectively. Threshold concentrations of IDPs are now predicted reasonably well by coarse-grained simulations where each amino acid is modeled by a single bead with a Lennard-Jones diameter *d*_0_ and a stickiness parameter *λ* (45). Our *q* parameter shows a good correlation (*R*^2^ = 0.59) with the compound parameter 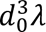 (Figure 9). Therefore the *q* parameter may serve as a predictor for the tendency of an amino acid to drive phase separation. In essence, the same ability of an amino acid, e.g., Trp, to form interactions with neighboring residues of an IDP in the free state also applies when it comes to interactions with residues on neighboring chains in a dense phase.

**Figure 9.**
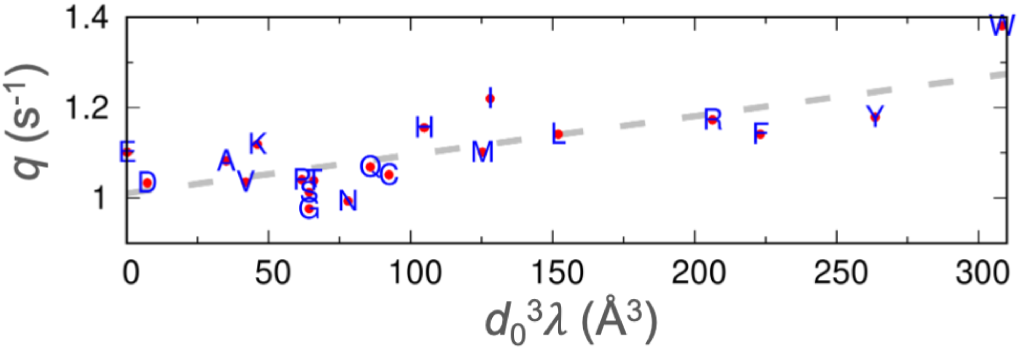
Correlation between the stickiness parameters (*λ*) and the NMR relaxation parameters (*q*). The regression line is shown as dashes.

Our method incorporates ideas from a number of previous efforts at describing *R*_2_. The first serious effort was by Schwalbe et al. (15), who accounted for contributions from neighboring residues as additive terms, instead of multiplicative factors as in SeqDYN. Cho et al. (16) and Delaforge et al. (24) used the running average of the bulkiness parameter over a window of five to nine residues as a qualitative indicator of *R*_2_. Here again the calculation was based on an additive model. Sekiyama et al. (17) employed a multiplicative model, with *R*_2_ calculated as a geometric mean of “indices of local dynamics” over a five-residue window. These indices, akin to our *q* parameters, were trained on a single IDR (TIA-1 prion-like domain) and used to reproduce the measured *R*_2_ for the same IDR. As we have illustrated on CBP-ID4 (Figure 5-figure supplement 1), training on a single protein merely biases the parameters to that model and has little value in predicting *R*_2_ for other proteins. In comparison, SeqDYN is trained on 45 IDPs and its predictions are robust and achieve quantitative agreement with measured *R*_2_.

Ten of the IDPs tested here have been studied recently by MD simulations using IDP-specific force fields (1, 12-14). In Table 3 we compare the RMSEs of SeqDYN predictions with those for *R*_2_ calculations from MD simulations. For five of these IDPs: A1-LCD, Aβ40, α-synuclein, tau K18, and FtsQ, RMSEs of SeqDYN and MD are remarkably similar. Four of these IDPs lack significant population of α-helices or β-sheets, but FtsQ forms a stable long helix. For one other IDP, namely HOX-DFD, MD, by explicitly modeling its folded domain, does a much better job in predicting *R*_2_ than SeqDYN (RMSEs of 1.40 s^-1^ vs 1.99 s^-1^). However, for the four remaining IDPs: p53TAD, Pup, Sev-NT, and ChiZ, SeqDYN significantly outperforms MD, with RMSEs averaging only 0.47 s^-1^, compared to the MD counterpart of 1.14 s^-1^. Overall, SeqDYN is very competitive against MD in predicting *R*_2_, but without the significant computational cost. While MD simulations can reveal details of local interactions, as noted for α-synuclein, and capture tertiary interactions if they occur, they still suffer from perennial problems of force-field imperfection and inadequate sampling. SeqDYN provides an accurate description of IDP dynamics at a “mean-field” level, but could miss idiosyncratic behaviors of specific local sequences.

**Table 3.** RMSEs (s^-1^) of *R*_2_ predictions by SeqDYN and MD for 10 IDPs.

| IDP name | SeqDYN | MD |
| --- | --- | --- |
| A1-LCD | 0.60<sup>a</sup> | 0.59<sup>d, e</sup> |
| Aβ40 | 0.38<sup>a</sup> | 0.38<sup>d</sup> |
| HOX-DFD | 1.99<sup>a</sup> | 1.40<sup>d</sup> |
| α-synuclein | 0.44<sup>a</sup> | 0.50<sup>d</sup> |
| p53TAD | 0.33<sup>a</sup> | 1.04<sup>f</sup> |
| Pup | 0.43<sup>a</sup> | 1.00<sup>f</sup> |
| Sev-NT | 0.38<sup>a, b</sup> | 1.10<sup>d, g</sup> |
| tau K18 | 0.83<sup>a</sup> | 0.80<sup>d</sup> |
| ChiZ | 0.74<sup>c</sup> | 1.40<sup>h</sup> |
| FtsQ | 1.71<sup>c, b</sup> | 1.70<sup>i</sup> |

<sup>a</sup>Based on leave-one-out training (using 44 IDPs).
<sup>b</sup>Helix boost applied.
<sup>c</sup>Based on training by the full training set (45 IDPs).
<sup>d</sup>From ref (1).
<sup>e</sup>RMSE is scaled down by a factor of 2.39, to correct for the effect of temperature (MD at 288 K; see Figure 3-figure supplement 1C).
<sup>f</sup>From ref (13).
<sup>g</sup>RMSE is scaled down by a factor of 2.99, to correct for the effects of temperature and magnetic field (MD at 274 K and 850 MHz; see Figure 3-figure supplement 1B).
<sup>h</sup>Originally calculated in ref (12) with correction in ref (80).
<sup>i</sup>From ref (14).

Deep-learning models have become very powerful, but they usually have millions of parameters and require millions of protein sequences for training (46). In contrast, SeqDYN employs a mathematical model with dozens of parameters and requires only dozens of proteins for training. Reduced models (by collapsing amino acids into a small number of distinct types) have even been trained on < 10 IDPs to predict propensities for binding nanoparticles (18) or membranes (19). The mathematical model-based approach may be useful in other applications where data, similar to *R*_2_, are limited, including predictions of IDP secondary chemical shifts or residues that bind drug molecules (47) or protein targets, or even in protein design, e.g., for recognizing an antigenic site or a specific DNA site.

## Methods

### Collection of IDPs with measured *R*_2_

Starting from six nonhomologous IDPs in our previous MD study (1), we obtained *R*_2_ data for eight IDPs from the Bimolecular Magnetic Resonance Data Bank (BMRB; https://bmrb.io); data for two other IDPs were from our collaborators (12, 14). Most of the 54 IDPs studied here were from searching the literature. Disorder was judged by dispersion in backbone amide proton chemical shifts, NOE, and SCS. *R*_2_ data that were not available from the authors or BMRB were obtained by digitizing *R*_2_ plots presented in figures of published papers, using WebPlotDigitizer (https://automeris.io/WebPlotDigitizer) (48) and further inspected visually.

Homology of IDPs was checked by sequence alignment using Clustal W (http://www.clustal.org/clustal2) (49), and presented as a clock-like tree using the “ape” package (http://ape-package.ird.fr) (50). IDPs that had discernible homology with the selected training set were removed. Removed IDPs included HOX-SCR and β-synuclein from our previous MD study (1), due to homology with HOX-DFD and α-synuclein, respectively.

### Coding for SeqDYN

The training of SeqDYN was coded in python, similar to our previous work for predicting residue-specific membrane association propensities (ReSMAP; https://zhougroup-uic.github.io/ReSMAPidp/) (19). The cost function was the sum of mean-squared-errors for the IDPs in the training set. We used the least_squares function in scipy.optimize, with Trust Region Reflective as the minimization algorithm and all parameters restricted to the positive range. For the web server (https://zhougroup-uic.github.io/SeqDYNidp/), we rewrote the prediction code javascript.

## ACKNOWLEDGEMENTS

This work was supported by Grant GM118091 from the National Institutes of Health.

## FIGURE CAPTIONS

**Figure 3-figure supplement 1.**
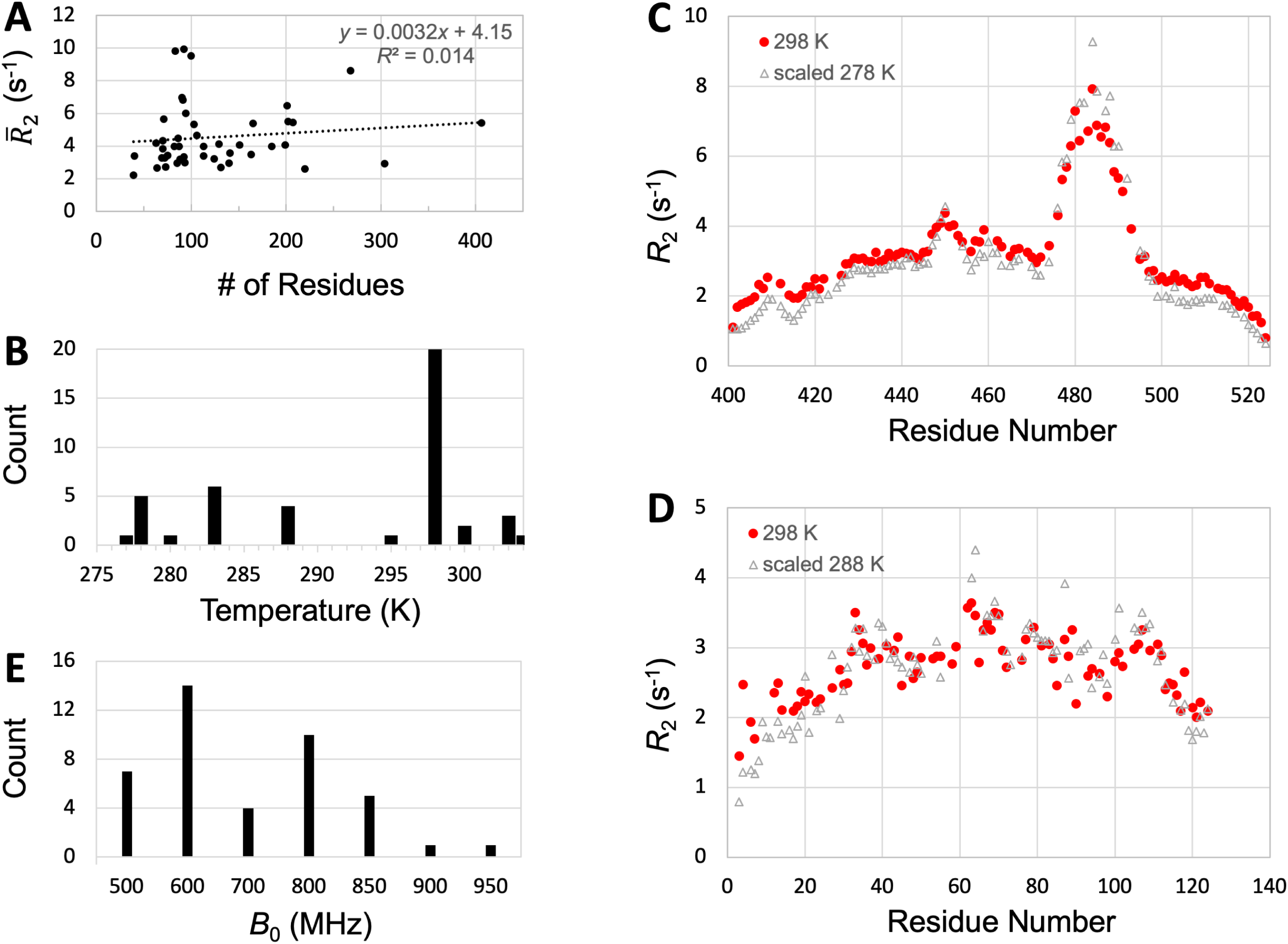
Possible effects of sequence length, temperature, and magnetic field on *R*_2_. (A) Lack of dependence of 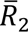 on sequence length. (B) Counts of IDPs with *R*_2_ measured at various temperatures. (C) Matching of *R*_2_ profiles of Sev-NT measured at two temperatures after uniform scaling. (D) Matching of *R*_2_ profiles of A1-LCD measured at two temperatures after uniform scaling. (E) Counts of IDPs with *R*_2_ measured at various magnetic fields.

**Figure 4-figure supplement 1.**
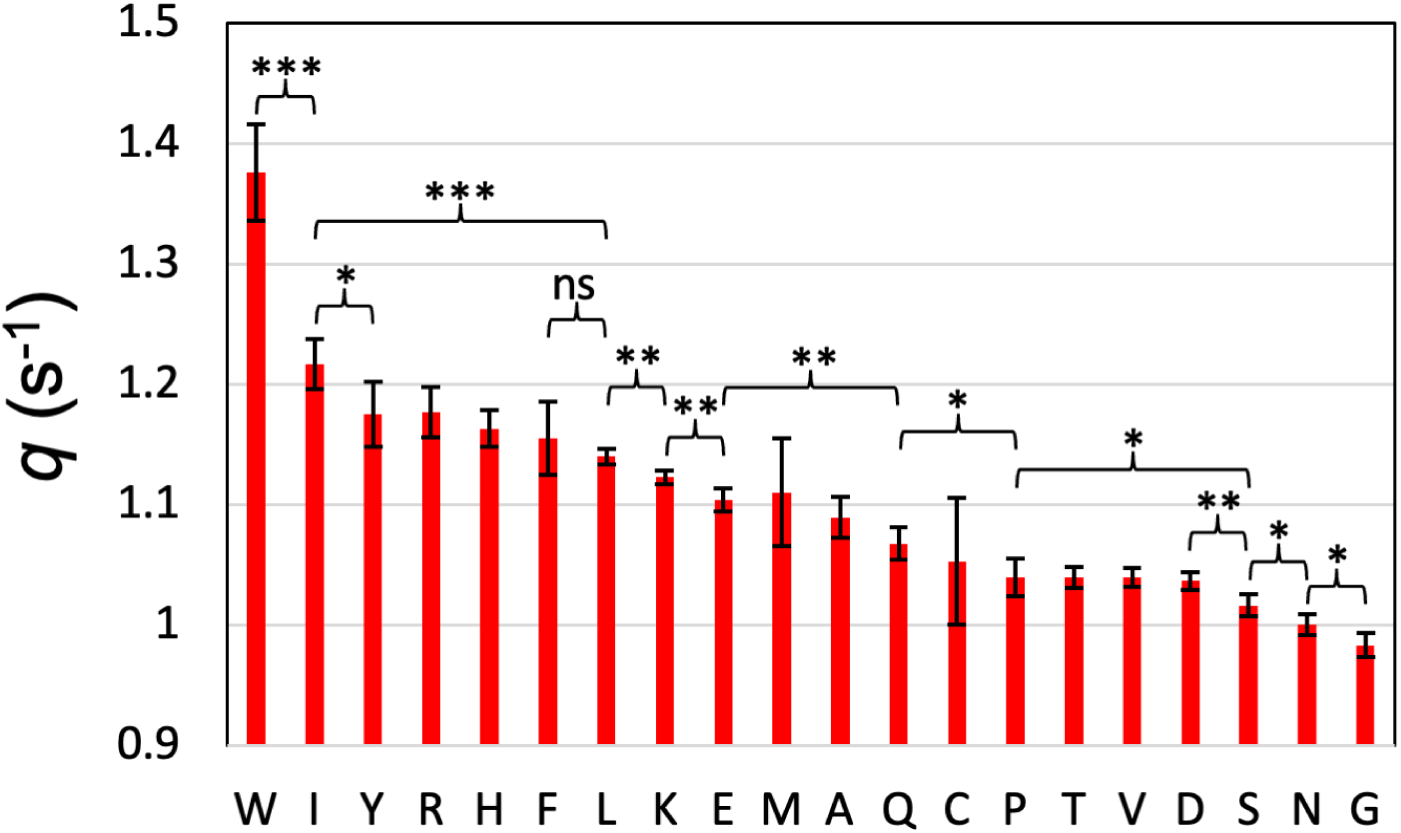
T-test of on the *q* parameters for pairs of amino acids. *q* parameters were obtained from five-fold cross-validation training, resulting in five independent values for each *q* parameter. Mean presented as red bars; standard deviation presented as error bar. *, *p* < 0.05; **, *p* < 0.01; ***, *p* < 0.001; ns, not significant. *q* parameters for all neighboring pairs not explicitly indicated are not significantly different.

**Figure 5-figure supplement 1.**
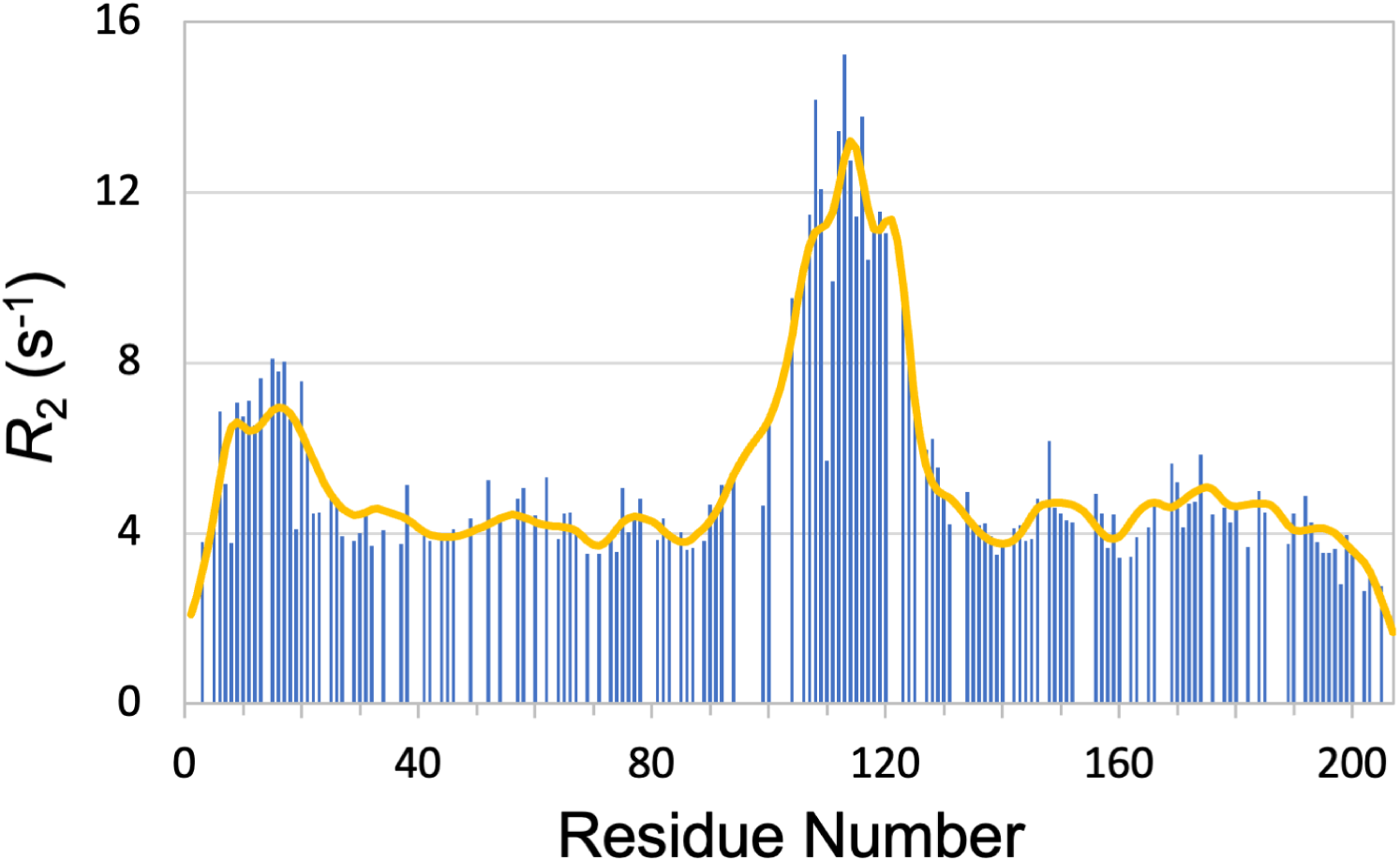
Close reproduction (curve) of the measured *R*_2_ profile (bars) of CBP-ID4 when that set of data alone was used to parameterize SeqDYN. The resulting model has no value for predicting *R*_2_ for other proteins.

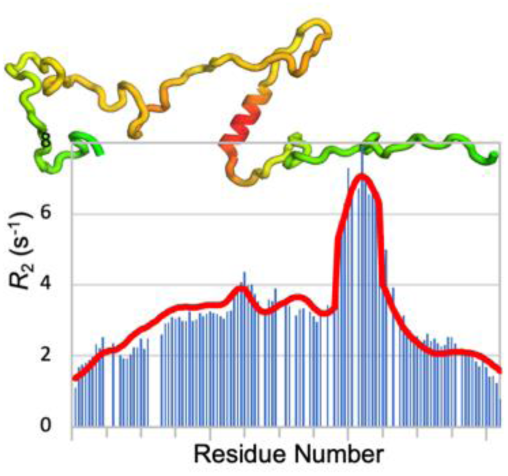

Dynamics is a crucial link between sequence and function for intrinsically disordered proteins (IDPs). Qin and Zhou developed a sequence-based method to predict IDP dynamics. The predictions provide deep physicochemical insight and can also serve as indicators of properties including phase-separation propensities of IDP sequences.

## Notes

### Competing Interest Statement

The authors have declared no competing interest.

### Summary of Updates

We moved all supplementary materials into the main manuscript.

## REFERENCES

1. S. Dey, M. MacAinsh, H. X. Zhou, Sequence-Dependent Backbone Dynamics of Intrinsically Disordered Proteins. J Chem Theory Comput 18, 6310–6323 (2022).

2. A. R. Camacho-Zarco et al., NMR Provides Unique Insight into the Functional Dynamics and Interactions of Intrinsically Disordered Proteins. Chem Rev 122, 9331–9356 (2022).

3. J. Klein-Seetharaman et al., Long-Range Interactions Within a Nonnative Protein. Science 295, 1719–1722 (2002).

4. A. Abyzov et al., Identification of Dynamic Modes in an Intrinsically Disordered Protein Using Temperature-Dependent NMR Relaxation. J Am Chem Soc 138, 6240–6251 (2016).

5. R. Schneider et al., Visualizing the Molecular Recognition Trajectory of an Intrinsically Disordered Protein Using Multinuclear Relaxation Dispersion NMR. J Am Chem Soc 137, 1220–1229 (2015).

6. M. Feichtinger et al., Long-range structural preformation in yes-associated protein precedes encounter complex formation with TEAD. iScience 25, 104099 (2022).

7. N.-A. Lakomek, H. Yavuz, R. Jahn, Á. Pérez-Lara, Structural dynamics and transient lipid binding of synaptobrevin-2 tune SNARE assembly and membrane fusion. Proc Natl Acad Sci U S A 116, 8699–8708 (2019).

8. A. E. Conicella, G. H. Zerze, J. Mittal, N. L. Fawzi, ALS Mutations Disrupt Phase Separation Mediated by alpha-Helical Structure in the TDP-43 Low-Complexity C-Terminal Domain. Structure 24, 1537–1549 (2016).

9. A. Malki et al., Intrinsically Disordered Tardigrade Proteins Self-Assemble into Fibrous Gels in Response to Environmental Stress. Angew Chem Int Ed Engl 61, e202109961 (2022).

10. E. W. Martin et al., Valence and patterning of aromatic residues determine the phase behavior of prion-like domains. Science 367, 694–699 (2020).

11. E. C. Cook, D. Sahu, M. Bastidas, S. A. Showalter, Solution Ensemble of the C-Terminal Domain from the Transcription Factor Pdx1 Resembles an Excluded Volume Polymer. J Phys Chem B 123, 106–116 (2019).

12. A. Hicks, C. A. Escobar, T. A. Cross, H. X. Zhou, Sequence-dependent correlated segments in the intrinsically disordered region of ChiZ. Biomolecules 10, 1–23 (2020).

13. L. Yu, R. Bruschweiler, Quantitative prediction of ensemble dynamics, shapes and contact propensities of intrinsically disordered proteins. PLoS Comput Biol 18, e1010036 (2022).

14. S. T. Smrt, C. A. Escobar, S. Dey, T. A. Cross, H. X. Zhou, An Arg/Ala-rich helix in the N-terminal region of M. tuberculosis FtsQ is a potential membrane anchor of the Z-ring. Commun Biol 6, 311 (2023).

15. H. Schwalbe et al., Structural and dynamical properties of a denatured protein. Heteronuclear 3D NMR experiments and theoretical simulations of lysozyme in 8 M urea. Biochemistry 36, 8977-8991 (1997).

16. M. K. Cho et al., Amino acid bulkiness defines the local conformations and dynamics of natively unfolded alpha-synuclein and tau. J Am Chem Soc 129, 3032–3033 (2007).

17. N. Sekiyama et al., ALS mutations in the TIA-1 prion-like domain trigger highly condensed pathogenic structures. Proc Natl Acad Sci U S A 119, e2122523119 (2022).

18. D. W. Li, M. Xie, R. Bruschweiler, Quantitative cooperative binding model for intrinsically disordered proteins interacting with nanomaterials. J Am Chem Soc 142, 10730–10738 (2020).

19. S. Qin, A. Hicks, S. Dey, R. Prasad, H. X. Zhou, ReSMAP: Web Server for Predicting Residue-Specific Membrane-Association Propensities of Intrinsically Disordered Proteins. Membranes 12, 773 (2022).

20. M. R. Jensen et al., Quantitative conformational analysis of partially folded proteins from residual dipolar couplings: application to the molecular recognition element of Sendai virus nucleoprotein. J Am Chem Soc 130, 8055–8061 (2008).

21. A. Piai et al., Just a Flexible Linker? The Structural and Dynamic Properties of CBP-ID4 Revealed by NMR Spectroscopy. Biophys J 110, 372-381 (2016).

22. S. Maiti et al., Dynamic Studies on Intrinsically Disordered Regions of Two Paralogous Transcription Factors Reveal Rigid Segments with Important Biological Functions. J Mol Biol 431, 1353–1369 (2019).

23. S. Patel, V. Ramanujam, A. K. Srivastava, K. V. Chary, Conformational propensities and dynamics of a betagamma-crystallin, an intrinsically disordered protein. Phys Chem Chem Phys 16, 12703–12718 (2014).

24. E. Delaforge et al., Deciphering the Dynamic Interaction Profile of an Intrinsically Disordered Protein by NMR Exchange Spectroscopy. J Am Chem Soc 140, 1148–1158 (2018).

25. Y. H. Sung, D. Eliezer, Residual structure, backbone dynamics, and interactions within the synuclein family. J Mol Biol 372, 689–707 (2007).

26. S. Milles et al., An ultraweak interaction in the intrinsically disordered replication machinery is essential for measles virus function. Sci Adv 4, eaat7778 (2018).

27. Z. Solyom et al., The Disordered Region of the HCV Protein NS5A: Conformational Dynamics, SH3 Binding, and Phosphorylation. Biophys J 109, 1483-1496 (2015).

28. E. W. Martin et al., Sequence Determinants of the Conformational Properties of an Intrinsically Disordered Protein Prior to and upon Multisite Phosphorylation. J Am Chem Soc 138, 15323–15335 (2016).

29. A. M. Janke et al., Lysines in the RNA Polymerase II C-Terminal Domain Contribute to TAF15 Fibril Recruitment. Biochemistry 57, 2549–2563 (2018).

30. M. Jenner et al., Mechanism of intersubunit ketosynthase-dehydratase interaction in polyketide synthases. Nat Chem Biol 14, 270–275 (2018).

31. E. M. Bafaro, M. W. Maciejewski, J. C. Hoch, R. E. Dempski, Concomitant disorder and high-affinity zinc binding in the human zinc- and iron-regulated transport protein 4 intracellular loop. Protein Sci 28, 868–880 (2019).

32. M. G. Murrali, A. Piai, W. Bermel, I. C. Felli, R. Pierattelli, Proline Fingerprint in Intrinsically Disordered Proteins. ChemBioChem 19, 1625–1629 (2018).

33. L. E. Wong, T. H. Kim, D. R. Muhandiram, J. D. Forman-Kay, L. E. Kay, NMR Experiments for Studies of Dilute and Condensed Protein Phases: Application to the Phase-Separating Protein CAPRIN1. J Am Chem Soc 142, 2471–2489 (2020).

34. J. M. Zimmerman, N. Eliezer, R. Simha, The characterization of amino acid sequences in proteins by statistical methods. J Theor Biol 21, 170–201 (1968).

35. L. J. McGuffin, K. Bryson, D. T. Jones, The PSIPRED protein structure prediction server. Bioinformatics 16, 404–405 (2000).

36. H. Y. Cho et al., Characterization of the interaction between lysyl-tRNA synthetase and laminin receptor by NMR. FEBS Lett 588, 2851–2858 (2014).

37. Y. Shen, F. Delaglio, G. Cornilescu, A. Bax, TALOS+: a hybrid method for predicting protein backbone torsion angles from NMR chemical shifts. J Biomol NMR 44, 213–223 (2009).

38. C. Thapa, P. Roivas, T. Haataja, P. Permi, U. Pentikainen, Interaction mechanism of endogenous PP2A inhibitor protein ENSA with PP2A. Febs J 289, 519–534 (2022).

39. A. Vogel, A. Crawford, A. Nyarko, Multivalent Angiomotin-like 1 and Yes-associated protein form a dynamic complex. Protein Sci 31, e4295 (2022).

40. S. Schwarzinger, P. E. Wright, H. J. Dyson, Molecular hinges in protein folding: the urea-denatured state of apomyoglobin. Biochemistry 41, 12681–12686 (2002).

41. J. Wirmer, W. Peti, H. Schwalbe, Motional properties of unfolded ubiquitin: a model for a random coil protein. J Biomol NMR 35, 175–186 (2006).

42. D. Ban, L. I. Iconaru, A. Ramanathan, J. Zuo, R. W. Kriwacki, A Small Molecule Causes a Population Shift in the Conformational Landscape of an Intrinsically Disordered Protein. J Am Chem Soc 139, 13692–13700 (2017).

43. J. Wang et al., A Molecular Grammar Governing the Driving Forces for Phase Separation of Prion-like RNA Binding Proteins. Cell 174, 688–699 e616 (2018).

44. Y. Zhang, R. Prasad, S. Su, D. Lee, H. X. Zhou, Amino Acid-Dependent Phase Equilibrium and Material Properties of Tetrapeptide Condensates. Cell Rep Phys Sci 5, 102218 (2024).

45. G. Tesei, K. Lindorff-Larsen, Improved predictions of phase behaviour of intrinsically disordered proteins by tuning the interaction range. Open Res Eur 2, 94 (2022).

46. A. Rives et al., Biological structure and function emerge from scaling unsupervised learning to 250 million protein sequences. Proc Natl Acad Sci U S A 118, e2016239118 (2021).

47. P. Robustelli et al., Molecular Basis of Small-Molecule Binding to alpha-Synuclein. J Am Chem Soc 144, 2501–2510 (2022).

48. A. Rohatgi, Webplotdigitizer: Version 4.6 (2022).

49. M. A. Larkin et al., Clustal W and Clustal X version 2.0. Bioinformatics 23, 2947–2948 (2007).

50. E. Paradis, J. Claude, K. Strimmer, APE: Analyses of Phylogenetics and Evolution in R language. Bioinformatics 20, 289–290 (2004).

51. N. Rezaei-Ghaleh, G. Parigi, M. Zweckstetter, Reorientational Dynamics of Amyloid-beta from NMR Spin Relaxation and Molecular Simulation. J Phys Chem Lett 10, 3369–3375 (2019).

52. S. Yao et al., Characterisation of the conformational preference and dynamics of the intrinsically disordered N-terminal region of Beclin 1 by NMR spectroscopy. Biochim Biophys Acta 1864, 1128–1137 (2016).

53. B. Szalaine Agoston, D. Kovacs, P. Tompa, A. Perczel, Full backbone assignment and dynamics of the intrinsically disordered dehydrin ERD14. Biomol NMR Assign 5, 189–193 (2011).

54. Z. Zheng, D. Ma, T. L. Yahr, L. Chen, The transiently ordered regions in intrinsically disordered ExsE are correlated with structural elements involved in chaperone binding. Biochem Biophys Res Commun 417, 129–134 (2012).

55. C. W. Lawrence, S. A. Showalter, Carbon-Detected (15)N NMR Spin Relaxation of an Intrinsically Disordered Protein: FCP1 Dynamics Unbound and in Complex with RAP74. J Phys Chem Lett 3, 1409–1413 (2012).

56. K. A. Burke, A. M. Janke, C. L. Rhine, N. L. Fawzi, Residue-by-Residue View of In Vitro FUS Granules that Bind the C-Terminal Domain of RNA Polymerase II. Mol Cell 60, 231–241 (2015).

57. T. Gruber et al., Macromolecular Crowding Induces a Binding Competent Transient Structure in Intrinsically Disordered Gab1. J Mol Biol 434, 167407 (2022).

58. M. O. Ebert, S. H. Bae, H. J. Dyson, P. E. Wright, NMR relaxation study of the complex formed between CBP and the activation domain of the nuclear hormone receptor coactivator ACTR. Biochemistry 47, 1299–1308 (2008).

59. R. Kiss, D. Kovacs, P. Tompa, A. Perczel, Local structural preferences of calpastatin, the intrinsically unstructured protein inhibitor of calpain. Biochemistry 47, 6936–6945 (2008).

60. F. C. Lopes et al., Pliable natural biocide: Jaburetox is an intrinsically disordered insecticidal and fungicidal polypeptide derived from jack bean urease. Febs J 282, 1043–1064 (2015).

61. M. De Avila, K. A. Vassall, G. S. Smith, V. V. Bamm, G. Harauz, The proline-rich region of 18.5 kDa myelin basic protein binds to the SH3-domain of Fyn tyrosine kinase with the aid of an upstream segment to form a dynamic complex in vitro. Biosci Rep 34, e00157 (2014).

62. S. Mokhtarzada, C. Yu, A. Brickenden, W. Y. Choy, Structural characterization of partially disordered human Chibby: insights into its function in the Wnt-signaling pathway. Biochemistry 50, 715–726 (2011).

63. M. Schiavina et al., Ensemble description of the intrinsically disordered N-terminal domain of the Nipah virus P/V protein from combined NMR and SAXS. Sci Rep 10, 19574 (2020).

64. J. L. Neira et al., Dynamics of the intrinsically disordered protein NUPR1 in isolation and in its fuzzy complexes with DNA and prothymosin alpha. Biochim Biophys Acta Proteins Proteom 1867, 140252 (2019).

65. B. Mateos et al., The Ambivalent Role of Proline Residues in an Intrinsically Disordered Protein: From Disorder Promoters to Compaction Facilitators. J Mol Biol 432, 3093–3111 (2020).

66. M. Xie, D. W. Li, J. Yuan, A. L. Hansen, R. Bruschweiler, Quantitative Binding Behavior of Intrinsically Disordered Proteins to Nanoparticle Surfaces at Individual Residue Level. Chemistry 24, 16997–17001 (2018).

67. J. Song et al., Intrinsically disordered gamma-subunit of cGMP phosphodiesterase encodes functionally relevant transient secondary and tertiary structure. Proc Natl Acad Sci U S A 105, 1505–1510 (2008).

68. C. Olivieri et al., Multi-state recognition pathway of the intrinsically disordered protein kinase inhibitor by protein kinase A. Elife 9 (2020).

69. A. Borgia et al., Extreme disorder in an ultrahigh-affinity protein complex. Nature 555, 61–66 (2018).

70. M. A. Ahmed, V. V. Bamm, G. Harauz, V. Ladizhansky, The BG21 isoform of Golli myelin basic protein is intrinsically disordered with a highly flexible amino-terminal domain. Biochemistry 46, 9700–9712 (2007).

71. V. Csizmok, I. C. Felli, P. Tompa, L. Banci, I. Bertini, Structural and dynamic characterization of intrinsically disordered human securin by NMR spectroscopy. J Am Chem Soc 130, 16873–16879 (2008).

72. T. Mittag et al., Structure/function implications in a dynamic complex of the intrinsically disordered Sic1 with the Cdc4 subunit of an SCF ubiquitin ligase. Structure 18, 494–506 (2010).

73. X. Wang et al., A large intrinsically disordered region in SKIP and its disorder-order transition induced by PPIL1 binding revealed by NMR. J Biol Chem 285, 4951–4963 (2010).

74. R. Thapar, G. A. Mueller, W. F. Marzluff, The N-terminal domain of the Drosophila histone mRNA binding protein, SLBP, is intrinsically disordered with nascent helical structure. Biochemistry 43, 9390–9400 (2004).

75. N. Rezaei-Ghaleh et al., Local and global dynamics in intrinsically disordered synuclein. Angew Chem Int Ed Engl 57, 15262–15266 (2018).

76. I. R. Chandrashekaran et al., Structure and Functional Characterization of the Conserved JAK Interaction Region in the Intrinsically Disordered N-Terminus of SOCS5. Biochemistry 54, 4672–4682 (2015).

77. P. Barre, D. Eliezer, Structural transitions in tau k18 on micelle binding suggest a hierarchy in the efficacy of individual microtubule-binding repeats in filament nucleation. Protein Sci 22, 1037–1048 (2013).

78. E. A. Cino, M. Karttunen, W. Y. Choy, Effects of molecular crowding on the dynamics of intrinsically disordered proteins. PLoS One 7, e49876 (2012).

79. J. Harris, M. Shadrina, C. Oliver, J. Vogel, A. Mittermaier, Concerted millisecond timescale dynamics in the intrinsically disordered carboxyl terminus of gamma-tubulin induced by mutation of a conserved tyrosine residue. Protein Sci 27, 531–545 (2018).

80. A. Hicks, M. MacAinsh, H. X. Zhou, Removing thermostat distortions of protein dynamics in constant-temperature molecular dynamics simulations. J Chem Theory Comput 17, 5920–5932 (2021).

81. H. J. Feldman, C. W. Hogue, Probabilistic sampling of protein conformations: new hope for brute force? Proteins 46, 8–23 (2002).

82. P. Bernado, M. Blackledge, A self-consistent description of the conformational behavior of chemically denatured proteins from NMR and small angle scattering. Biophys J 97, 2839–2845 (2009).

